# A male seminal fluid protein SFP-1 regulates hermaphrodite post-mating longevity and fat metabolism in *Caenorhabditis elegans*

**DOI:** 10.1101/2024.09.09.612157

**Authors:** Mingqing Chen, Jianke Gong

**Affiliations:** Key Laboratory of Molecular Biophysics of MOE, and College of Life Science and Technology, Huazhong University of Science and Technology, Wuhan, Hubei 430074, China

## Abstract

There are several causes of mating-induced physiological changes in hermaphrodites exposed to males, such as functional male sperms causing shrinking and the male pheromone mediating shortened lifespan, which utilize different molecular pathways and are shared across species. However, it is unclear whether the male seminal fluid protein contributes to this post-mating regulation. Here, we investigated the transmit way and the impacts of the *Caenorhabditis* male seminal fluid protein SFP-1 in mated hermaphrodite tissues. We find that SFP-1 is the key component in seminal fluid to induce post-mating physiological changes in mated hermaphrodites. It acts as a cargo packing into exophers which require the phospholipid scramblase ANOH-1 and ANOH-2 to develop in male seminal vesicle. Exophers carrying SFP-1 cross over the somatic gonad uterus and eventually the protein is uptaken by the intestinal cells via endocytosis. Within the intestine, the NTF2-like domain of SFP-1 assists the association and interaction with the transcription factors SKN-1 and DAF-16 to induce post-mating somatic fat depletion and a shortened lifespan. Our study reveals the elaborate strategies of the male seminal fluid protein on triggering mating-induced physiological changes elicited by sexual interactions that could exist in other species.

## Introduction

Many species have both male and female individuals and fertilize the next generation through sexual coexistence and the combination of the two types of gametes. The interactions between the sexes are common in animals across evolutionary distances and can significantly influence the development, metabolism, and aging of both sexes ^1–5^. In mice, even vasectomized males cause the co-house female to gain more body weight and shorten the lifespan, the same effects are observed when the males are intact ^6^. In *Drosophila*, males downregulate the post-mated aging of females by sex peptides in seminal fluid ^7^. However, the underlying mechanisms of sex interaction affecting metabolism and aging are still unclear.

*Caenorhabditis elegans* can either self-fertilize as a hermaphrodite or cross-fertilize by mating with males ^8^. Under normal conditions, *C. elegans* populations are not 50:50 mixed. In laboratories, hermaphrodites are the dominant members cultivated with males being extremely rare (0.2%) ^9,10^. However, due to food scarcity or other factors, the proportion of male nematodes in natural habitats, especially in low nutrient circumstances, might greatly increase compared with cultured in laboratory ^9,10^. The cross-fertilization results in 50% male production, which also contributes to the amplification of the male population. Therefore, the sexes coexistence happens quite often in the wild. Although hermaphrodites can enhance their fertility and brood size by mating with males, the sperm and seminal fluid from males transported into hermaphrodites through the mating process induce post-mating responses ^11^. These male components elicit gene expression, metabolic processes, and many other physiological changes in hermaphrodites. Through a DAF-9/DAF-12 dependent pathway, the spermatozoa can cause the hermaphrodite shrinking after mating ^11^. Moreover, many metabolic genes, including the acyl-CoA binding protein ACBP-3, are up-regulated in mated hermaphrodites via sperm ^12^. The seminal fluid components that are transferred into hermaphrodites can disrupt the nuclear localization of the transcription factor FOXO/DAF-16, which is a classical lifespan regulation pathway reinforced by INS-7 feedback ^11^. Additionally, these components can influence a group of neuron-enriched genes such as *delm-2* that control the metabolism of intestinal fat ^12^. However, the specific components or factors in male seminal fluid affecting hermaphroditic metabolism and lifespan regulation after mating are still mostly unknown.

Seminal fluid contains a great variety of components, such as sugars, peptides, inorganic salts, small organic molecules, and proteins, which are mainly produced by sex accessory glands. Particularly, the proteins secreted by the seminal vesicle, vas deferens, or other accessory glands perform many physiological functions including sperm motility and capacitation regulation and the modulation of immune function ^13–15^. Several proteins, for instance, drosomycin, an antifungal protein, derived from the ejaculatory ducts, are transferred to females with seminal fluid during copulation to induce immunostimulatory properties in *Drosophila* ^16^. Additionally, a variety of fly proteins, like sex peptide proteins, act as the factors to remote female reproductive behavior and change female post-mating energy balance to enhance egg production ^17^. The above two examples illustrate the effects on females. The other one of the important functions of these proteins is to play a role in the maturation and activation of sperm. Heparin-binding proteins, which stimulate sperm capacitation in the female reproductive tract, are secreted into seminal fluid by male accessory glands in bulls and rats ^18^. In addition, bovine seminal fluid also contains a variety of bovine seminal plasma proteins modulating spermatozoa maturation through their calmodulin-binding activity ^19^. Similarly, the proteins in the seminal fluid of the nematode also have these conserved functions. In *C. elegans* males, TRY-5, a seminal fluid protease, functions as an extracellular activator to regulate sperm activation ^20^. Besides modulating female physiological processes and male sperm maturation, most of the proteins in seminal fluid still have many covered roles that have not been studied well.

Here, we reported the identification of a male protein, SFP-1 (F56D2.8), which was expressed in the seminal vesicle, secreted into seminal fluid, and had the properties expected of a hermaphrodite-biased regulator in post-mating lifespan and lipid metabolism. Loss of *sfp-1* in males suppressed the mating-induced shortened lifespan in mated hermaphrodites. Furthermore, before seminal fluid ejaculation, SFP-1 packing into the exophers required the mammalian phospholipid scramblase TMEM16F homolog ANOH-1 and ANOH-2 in male seminal vesicle and then cross over the somatic gonad uterus and eventually spread into the intestinal cells via endocytosis. Within the intestine, the NTF2-like domain assisted SFP-1 in associating with the transcription factors SKN-1 and DAF-16 to induce post-mating somatic fat loss and shorten the lifespan. In summary, our findings suggest that SFP-1 is the first protein demonstrated to be a transferred component in the seminal fluid into *C. elegans* hermaphrodite during the sex interaction, targeting the intestine cells to play a vital role in the regulation of post-mating longevity and lipid metabolism.

## Results

### SFP-1 is a seminal fluid protein and up-regulated by sex interaction to promote hermaphrodite post-mating short lifespan

The expression level of some proteins in male seminal fluid is regulated by the response to mating with hermaphrodites, although these proteins have not been investigated. Genes encoding secreted proteins F56D2.8, K12H6.5, C16C8.10, and F40G9.15 from seminal vesicle are up-regulated according to genome-wide transcriptional studies of mated and unmated males ^21,22^ (**Figure 1A**). These four proteins are all in small size and have simple three-dimensional structures. Meanwhile, there is a significant down-regulation of *grd-4*, which codes for the hedgehog-like domain protein produced from the male tail ^23^. It is still unknown why these genes express differently after mating. They may contribute to both male and hermaphrodite reproduction and post-mating physiological regulation. Therefore, the protein of unknown function in male seminal fluid has the potential to be an interactive factor for regulating the metabolism and death of hermaphrodites after mating. F56D2.8 is one of the up-regulated male-specific proteins and only expressed in seminal vesicle by transcriptomic assay in cryo-sectioned adult male worms ^22^. Therefore, we named the gene that encodes this secreted protein as *<u>s</u>eminal fluid <u>p</u>rotein 1*, *sfp-1*. SFP-1 has 133 amino acids containing not only signal peptides in the N terminal, but especially a nuclear transport factor 2 (NTF2) like domain which plays an important role in the trafficking of macromolecules, ions, and small molecules between the cytoplasm and nucleus, through the predictive fold assay ^24,25^ (**Figure 1B**). Structurally, this protein comprises four beta-sheets, three alpha helices, and five loops (**Figure 1C**). SFP-1 has the potential to act as an assistant for protein trafficking in or out from the cytoplasm to the nucleus.

**Figure 1:**
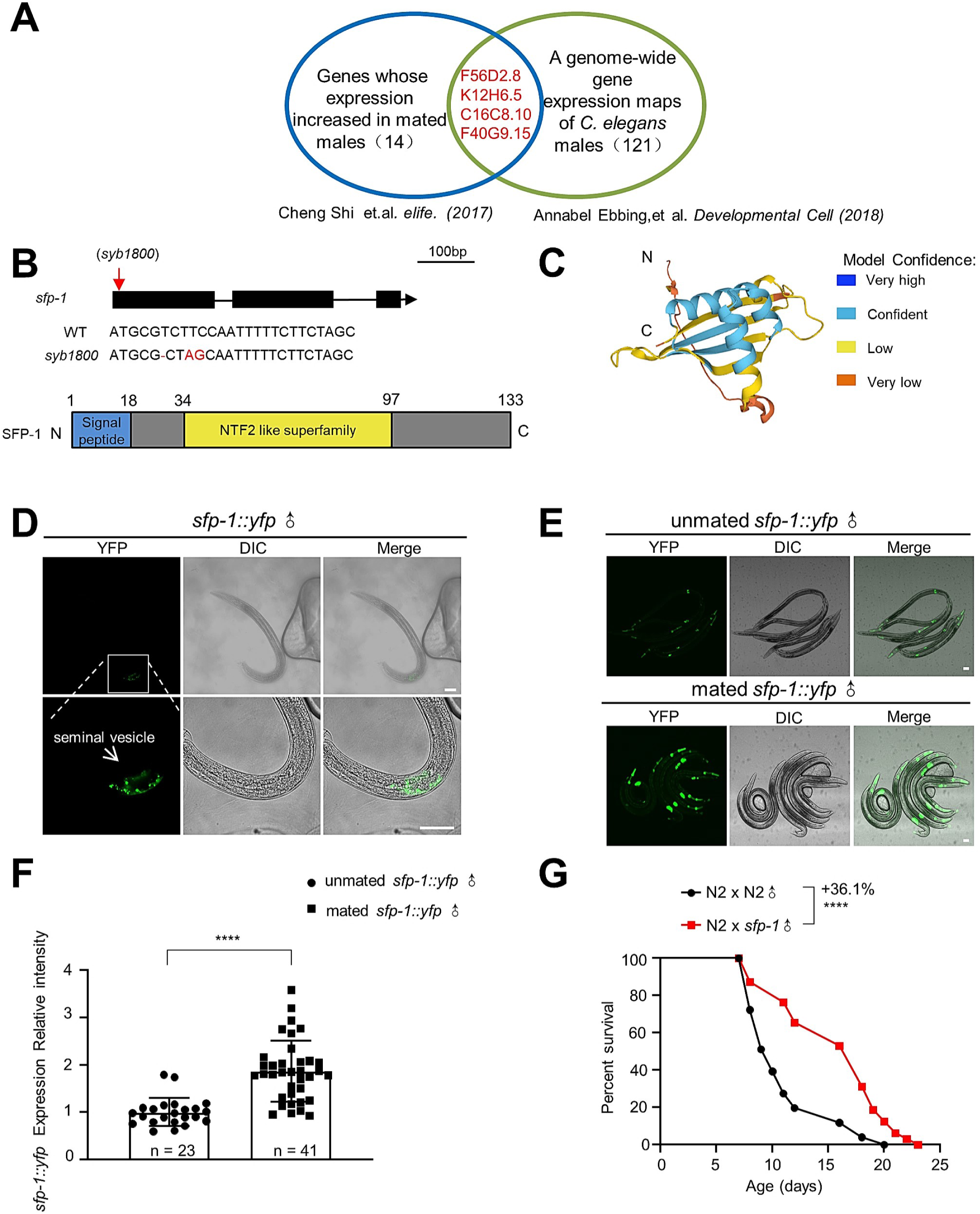
The secreted protein SFP-1 is involved in male mating and regulating longevity of mated hermaphrodites. (A) Genes that are upregulated in mated males and shared in male-specific genes. (B) Cartoon of genomic location of sfp-1 mutations. Domain architecture of SFP-1 and fragments, including signal peptide and NTF2L. (C) The AlphaFold2-Multimer model was used to predict the SFP-1 structure. (D) *sfp-1::yfp* is present in the somatic part of the male reproductive tract in the seminal vesicle. (Scale bars: 50 μm, 15 μm) (E-F) *sfp-1::yfp* expression increased in mated males. Mating on day1, pictures were taken on Day 2 of adulthood. Representative images are shown (E), the quantification of *sfp-1::yfp* expression (F).\*\*\*\**P* < 0.0001, t-test. (G) Lifespan of mated N2 worms. N2 x N2 ♂: 11.01 ± 0.66 days, n = 30 worms; N2 x *sfp-1*♂: 14.98 ± 0.72 days, n = 30 worms, *P* ˂ 0.001. Kaplan-Meier analysis with log-rank (Mantel-Cox) method was performed to compare the lifespans of different groups in this study. See Supplemental Table 1 for all lifespan data summary.

Even though a transcriptome-wide gene expression pattern analysis suggested that *sfp-1* is a male-specialized gene ^22^, its gene expression pattern has not been verified using fluorescent reporters. We generated a *sfp-1*::*yfp* fusion protein transgene line using the *sfp-1* gene its own promoter and performed confocal imaging. No fluorescence signal was detected in the adult hermaphrodite germline or sperm. However, there was a strong fluorescent signal in the seminal vesicle of males (**Figure 1D**), which indicated that SFP-1 is specifically expressed there and consequently resulted in a seminal fluid protein. Additionally, we noticed that in the seminal vesicle, the mated males of the transgenic line showed a higher fluorescent signal than the unmated males (**Figure 1E and 1F**). These findings reveal that SFP-1 might play an important role in sex interaction by acting as a seminal fluid factor mediating post-mating effects. Thus, we used CRISPR-cas9 technology to generate a loss-of-function allele of the *sfp-1* gene (**Figure 1B**). We performed mating assays by mating the wild-type hermaphrodites with either WT males or *sfp-1* knockout males and found that the lifespan of post-mating hermaphrodites mated with *sfp-1* mutants showed a prolonged lifespan (**Figure 1G**). However, compared with the WT mating group, the hermaphrodite brood size was enhanced and sperm-dependent post-mating shrinking was still triggered by the *sfp-1* mutant male (**Figure S1A and S1B**). Moreover, the *sfp-1* mutant male-conditioned plates were still effective in initiating the short lifespan in cultured hermaphrodites (**Figure S1C**). These results imply that SPF-1 does not affect sperm function and pheromone production ^26^. Furthermore, the over-expressed SFP-1 with its own promotor, *sfp-1::yfp,* male animals further shortened the mated hermaphrodites lifespan(**Figure S1D**). Thus, SFP-1 is the one of components in seminal fluid, turns on a high-expression pattern during sex interaction, and regulates hermaphrodite longevity after mating via the sperm and pheromone independent strategy.

### SFP-1 spreads into the intestine cells from the uterus in mated hermaphrodites via endocytosis

Seminal fluid was ejaculated into the uterus tissue of the hermaphrodite somatic gonad from the vulva after mating with males. SFP-1 should be able to fill up the uterus region nearby the vulva and then might be transported to other tissues. To test this hypothesis, we used the *glo-4* mutant hermaphrodite, which encodes a guanine nucleotide exchange factor required for biogenesis of the lysosome-related gut granules ^27^. This strain leads to better detection of weak YFP signals as the autofluorescence in the intestinal cells is much weaker than the wild-type animals ^28^. We performed confocal imaging for the *glo-4* mutant hermaphrodites mated with males expressing the *sfp-1::yfp*. We observed strong YFP signals in the uterus region near the vulva when the mating was just finished. Surprisingly, the fluorescence signals were detected in the intestine cells after 0.5 hour post-mating (**Figure 2A**). In contrast, no fluorescence signal in the intestine of *glo-4* mutant hermaphrodite mated with the wild-type males (**Figure 2A**). We imaged multiple mated hermaphrodites and observed similar signals in the intestinal cells (**Figure S2A**). To further demonstrate the transport capacity of SFP-1, we used MitoTracker to label the strain *sfp-1::yfp* male sperm. We observed rare translocated signals of the protein SFP-1 and sperms in the mated animal uterus (**Figure S2B**). Thus, SFP-1 requires another approach instead of the sperm to spread into gut cells. Moreover, for testing the endocytosis is the approach of the SFP-1 transport, we applied the endocytosis inhibiting drug, Bafilomycin A1 (bafA1), which prevents acidification-dependent vesicle turnover required for endosome maturation ^29^. During the mating processing, the mated hermaphrodites with bafA1 treatment had longer lifespan compared with the control mated animals (**Figure 2B**). We also performed the post-mating lifespan assay in the intestinal specific RNAi with the endocytosis key genes, such as *cav-1*, and *rab-10* ^30–32^. We found that the RNAi hermaphrodites when mated with males had significant longer post-mating lifespan than control (**Figure 2C-D**). According to these findings, SFP-1 within the seminal fluid is ejaculated into the mated hermaphrodite uterus region and then spread over the intestinal cells by endocytosis.

**Figure 2:**
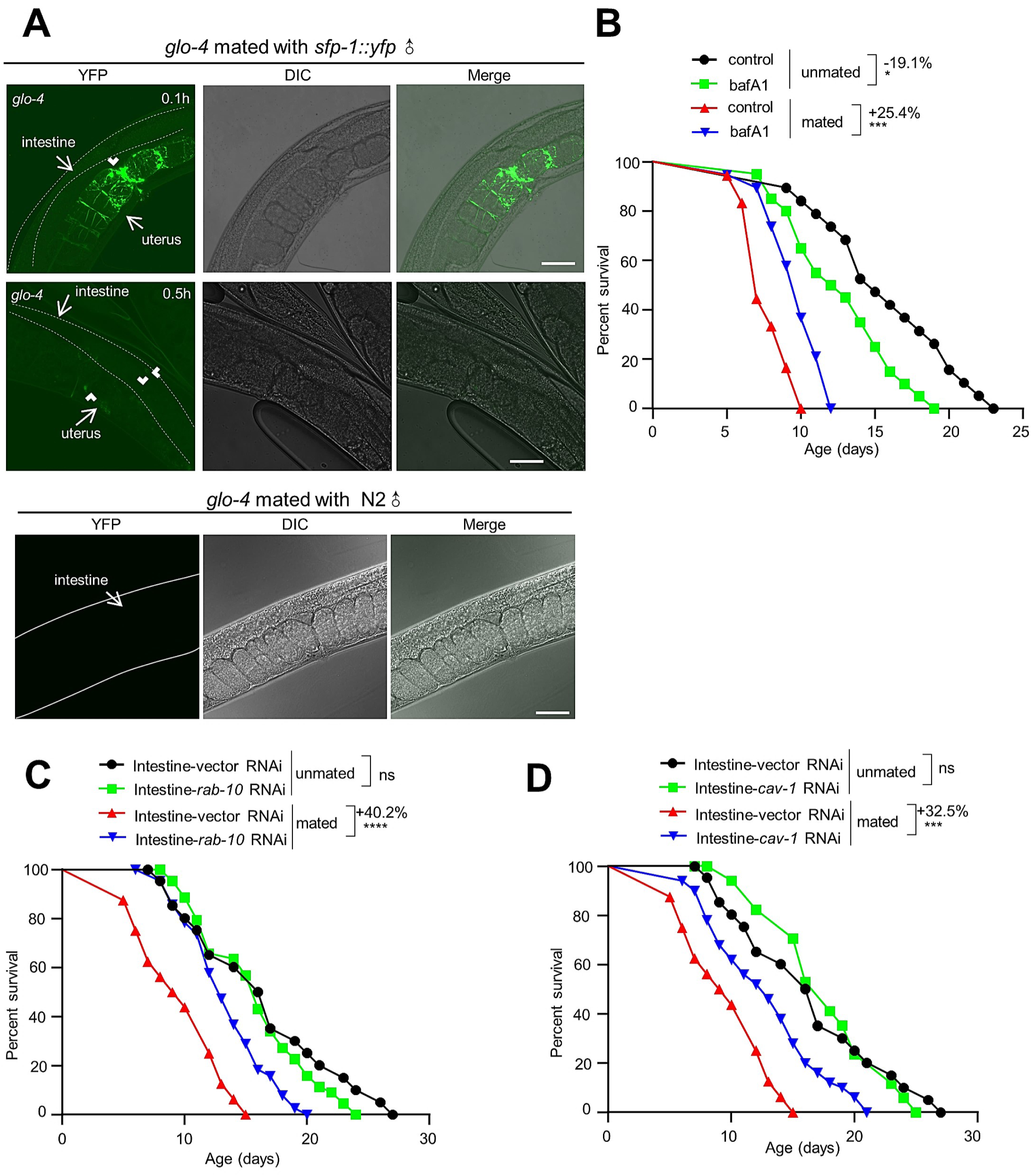
SFP-1 spreads into the intestine cells from the uterus in mated hermaphrodites via endocytosis. (A) Male-to-hermaphrodite transfer experiment where *sfp-1::yfp* males were mated with low autofluorescence *glo-4* hermaphrodites. Imaging of the low autofluorescence *glo-4* hermaphrodite after copulation revealed the presence of male derived *sfp-1::yfp* in the uterus, and then half an hour after mating, lots of weak fluorescence appeared in the intestine cell. No fluorescence signal was observed in the intestine cells of mated *glo-4* mutant. (B) Lifespan of mated and unmated N2 worms treated with the drug. N2 treated with DMSO: 15.63 ± 0.97 days, n = 19 worms; N2 treated with bafA1: 12.65 ± 0.79 days, n = 20 worms; N2 x N2 ♂ treated with DMSO: 7.72 ± 0.35 days, n = 20 worms; N2 x N2 ♂ treated with bafA 1: 9.68 ± 0.44 days, n = 21 worms, \*\**P* < 0.01,\*\*\**P* < 0.001. (C) Lifespan of mated and unmated intestine-specific *rab-10* RNAi. VP303-vector RNAi-unmated: 17.40 ± 0.31 days, n = 20 worms; VP303-*rab-10* RNAi-unmated: 15.84 ± 0.65 days, n = 44 worms; VP303-vector RNAi-mated: 9.63 ± 0.85 days, n = 16 worms; VP303-*rab-10* RNAi-mated: 13.49 ± 0.51 days, n = 39 worms, \*\*\*\**P* < 0.0001. (D) Lifespan of mated and unmated intestine-specific *cav-1* RNAi. VP303-vector RNAi-unmated: 17.40 ± 0.31 days, n = 20 worms; VP303-*cav-1* RNAi-unmated: 17.76 ± 1.06days, n = 17 worms; VP303-vector RNAi-mated: 9.63 ± 0.85 days, n =16 worms; VP303-*cav-1* RNAi-mated: 12.76 ± 0.62 days, n = 50 worms, \*\*\**P* < 0.001.

### Transportation of SFP-1 requires the phospholipid scramblase ANOH-1/ANOH-2 in males

We observed SFP-1 spread into the intestinal cells, however, how does it cross the border in the uterus. Very few macromolecules such as proteins can go through tissues by passive diffusion way. As a result, SFP-1 may be transported as cargo inside a bilayer vesicle of a male seminal vesicle. The lipid bilayer formed into large vesicles, known as exophers, which contain proteins, RNAs, lipids, and metabolites, mediating the intercellular and inter-tissue transport of biological macromolecules even organelles in most if not all cell types ^33,34^. The production of exophers from muscle cells is regulated by male pheromones, as reported recently ^35^. Lipid asymmetry between the inner and outer leaflets of the plasma membrane induces curvature to drive lipid bilayer vesicles release ^36,37^. Many enzymes are vital for the maintenance of bilayer asymmetry. Phospholipid scramblases ANOH-1/ANOH-2 translocate phospholipids from the inner to the outer leaflet and more importantly trigger the phosphatidylserine facilitated membrane fusion ^38–41^. Phospholipid scramblase mutants disrupt membrane asymmetry and fusion, thereby defecting the macromolecule transport because of the failure of the exophers formation and fusion in worms. According to confocal images of the *sfp-1::yfp* transgenic line, we observed that mating increased the amount of SFP-1 secreted into the exophers as cargo from the seminal vesicle. We observed many fluorescence exopher particles in the seminal vesicle based on the zoomed-in images (**Figure 3A**). However, compared to the wildtype male, there were significantly fewer exophers carrying SFP-1 in the single mutant background males, *anoh-1* and *anoh-2* (**Figure 3A**). In the seminal vesicle segment of the double mutant *anoh-1;anoh-2* male, SFP-1 was found in a diffuse state with the least amount of exophers (**Figure 3B**). Meanwhile, the double mutant males preeminently decreased the hermaphrodite post-mating death (**Figure 3C**). This phenotype did not show in the single mutant males, which may be due to the dose effect of the remaining exophers (**Figure 3C**). According to these findings, SFP-1 is transported from the seminal vesicle into exophers, which then use membrane fusion to transfer SFP-1 from the uterus to various tissues, primarily the intestine, to shorten the lifespan of post-mating hermaphrodites.

**Figure 3:**
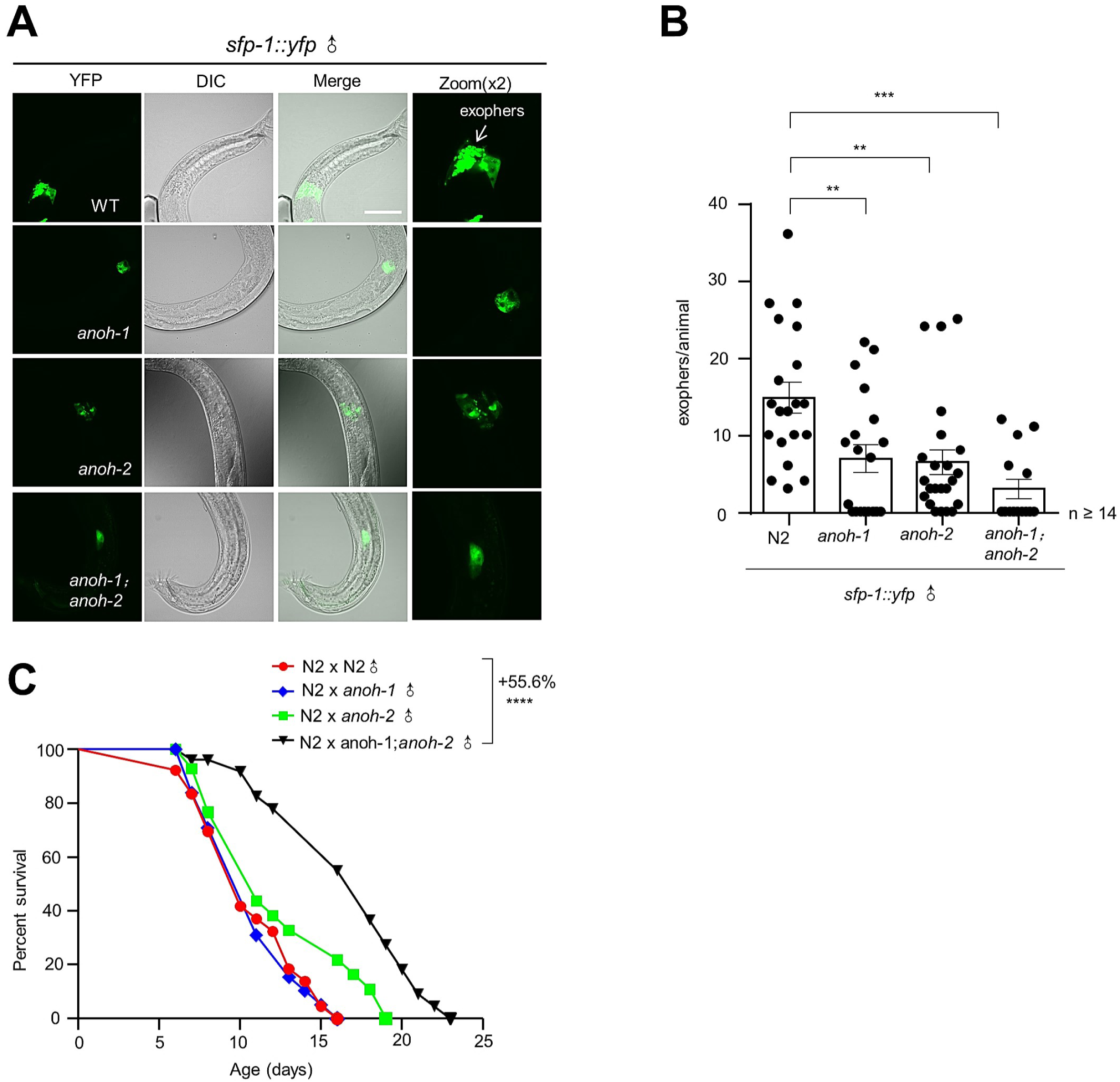
Transportation of SFP-1 requires phospholipid scramblase ANOH-1/ANOH-2 in males. (A) A *C. elegans* seminal vesicle exophers, Representative images of *sfp-1::yfp* in wildtype background and *anoh* single mutant and double mutant background. (Scale bars: 15 μm) (B) Lack of ANOH leads to a lower exopher genesis level in males. n = 20, 19, 23, and 14 worms (for respective columns). \*\**P* < 0.01,\*\*\**P* < 0.001 and *P* values were obtained by one-way ANOVA, Bonferroni’s multiple comparisons test. (C) Lifespan of mated N2 worms. N2 x N2 ♂: 10.45 ± 0.64 days, n = 22 worms; N2 x *anoh-1* ♂: 10.52 ± 0.56 days, n = 23 worms; N2 x *anoh-2* ♂: 11.93 ± 0.93 days, n = 20 worms; N2 x *anoh-1;anoh-2* ♂: 16.26 ± 0.91 days, n = 22 worms, \*\*\*\**P* < 0.0001. (Compared to hermaphrodite mated with N2 male).

### Ectopic expression of SFP-1 in the intestine cells imitates the post-mating phenotypes

We found that SPF-1 was transported into intestinal cells then it has a certain chance that SFP-1 may function as a phenotype trigger factor in the mated hermaphrodite intestine. To address this hypothesis, we ectopically expressed SFP-1 full-size protein to four major tissues including intestine, germline, muscle, and neuron by their specific promoters. Only the ectopic expression of SFP-1 in intestine cells shortened the lifespan (**Figure 4A**), interestingly, we also found that SFP-1 in the gut enhanced reproduction activity increasing the brood size of the transgenic animal (**Figure S1A**). Meanwhile, SFP-1 induced the shrinking phenotype through the intestine cell in the transgenic animal even without mating (**Figure S1B**), indicating that SFP-1 may activate the germline in the gut signal pathway to induce shrinking. Moreover, we did not find any protein leaking to other tissues when we ectopically expressed full-length SFP-1 in the intestine or muscle (**Figure S3B**). This suggests that the secretory signal peptide only functions in the seminal vesicle and the ectopic expression approach is demonstrated to have no side effects. We ectopically expressed the other seminal fluid proteins such as K12H6.5 and F40G9.15 in the intestine and no shortened lifespan phenotype was observed (**Figure S3A**). These findings imply that SFP-1 functions in the intestine cells mediating the post-mating phenotype and the ectopic expression approach of this secreted protein is effective and credible.

**Figure 4:**
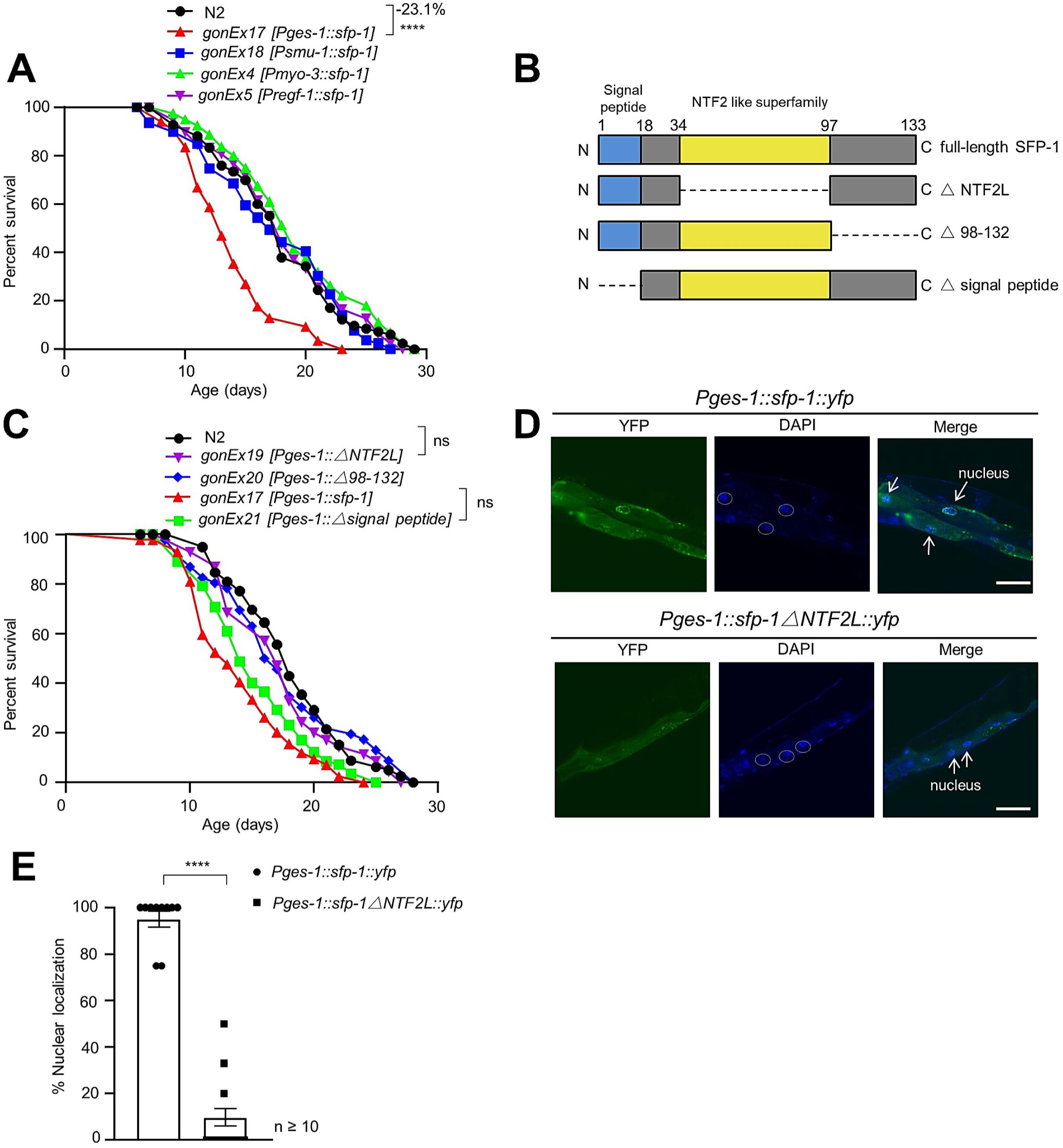
Only intestine over-expression SFP-1 can shorten the transgene line lifespan. (A) Lifespan survival curves of WT animals and tissue-specific expression SFP-1 in *Pges-1* (intestine), *Pmyo-3* (muscle), or *Prgef-1* (pan-neuronal) animals. Only the ectopic expression of SFP-1 in intestine cells shortened the lifespan. (B) Domain architecture of SFP-1 and fragments, including signal peptide and NTF2L. Δ, deletion. (C) Lifespan survival curves of WT animals and truncation strains. Lack of signal peptide in SFP-1 was unable to change the shortened lifespan. Compared to the control, the longevity difference was abolished in the absence of NTFL. (D-E) Representative pictures of nuclear localization in Day 1 *Pges-1::sfp-1::yfp* and *Pges-1::sfp-1△NTF2L::yfp* animals. (Scale bars: 15 μm). (E) Quantification of the nuclear localization of intestinal cells per individual. n > 10. \*\*\*\**P* < 0.0001 using t-test.

### NTF2-like domain is required for SFP-1 nuclear localization which causes post-mating like phenotype

According to the blast analysis, nearly half of the full-length amino acid sequence of SFP-1 forms a nuclear transport factor 2 like domain (NTF2L), therefore SFP-1 belongs to the NTF2-like superfamily. The NTF2-like superfamily comprises a large number of bacteria, archaea, and eukaryotes ^42,43^. This domain functions as a mediator of enzyme catalysis and protein-protein interaction evolutionally within the nuclear ^24^. To determine the function of SFP-1, we established three different truncations and found that the versions lacking NTF2L and C-terminal fragment (98-132 aa) eliminated the function of SFP-1 in mediating shortened lifespan in the ectopic expression worm (**Figure 4B and 4C**). Besides, the versions of lacking NTF2L protein failed to induce fat loss and enhance reproduction activity which we observed on the full-length protein version in the transgenic animals (**Figure S3C-S3E**). Furthermore, we observed that the full-length SFP-1 located in the gut cells nuclear where the NTF2L family protein presented its function using the *sfp-1::yfp* fusion protein transgenic line, but not the NTF2L truncation which SFP-1 remained in the cytoplasm (**Figure 4D and 4E**). Therefore, our results suggest that NTF2 like domain is essential for the SFP-1 translocation into nuclear and leading to the post-mating like phenotypes.

### SFP-1 is associated with the translocation of the FOXO transcription factor homolog DAF-16 after mating

Previous studies reported that mating induced DAF-16::GFP translocation out of the nucleus to counteract its promoting long lifespan genes activity via seminal fluid in *daf-2* hermaphrodites, while the component mediating this effect is still unknown ^44^. Whether SFP-1 plays a critical role in the association with DAF-16 to leave the nucleus requires to be explored. We used the protein interaction docking analysis website to anticipate the detailed features of the interfaces between the SFP-1 and DAF-16 to test this hypothesis ^44^. We discovered that those two proteins were held together directly by hydrogen bonds created by three residues in the contact surface (**Figure S4E**). Furthermore, we used *sfp-1* mutant male to mate with *daf-2* RNAi hermaphrodite in DAF-16::GFP expression line. Compared with the wildtype male which caused significant cytoplasmic localization of DAF-16, we observed most DAF-16::GFP fluorescence signals remained in the nucleus in the hermaphrodite mated with *sfp-1* mutant male (**Figure 5B**). Moreover, *sfp-1* mutant male rescued the mating-induced death in *daf-16* mutant background hermaphrodite, compared with the wildtype male which had an enhanced shorter lifespan (**Figure 5A**). Instead of having no effect on prolonging shortened lifespan phenotype, this indicates that SFP-1 induces post-mating aging via multiple pathways, maybe involving fat loss dependent lifespan shortening, not only the DAF-16 transcription factor dependent pathway. These data show that SFP-1 assists DAF-16 translocation out of the nucleus to partially promote mating-induced death.

**Figure 5:**
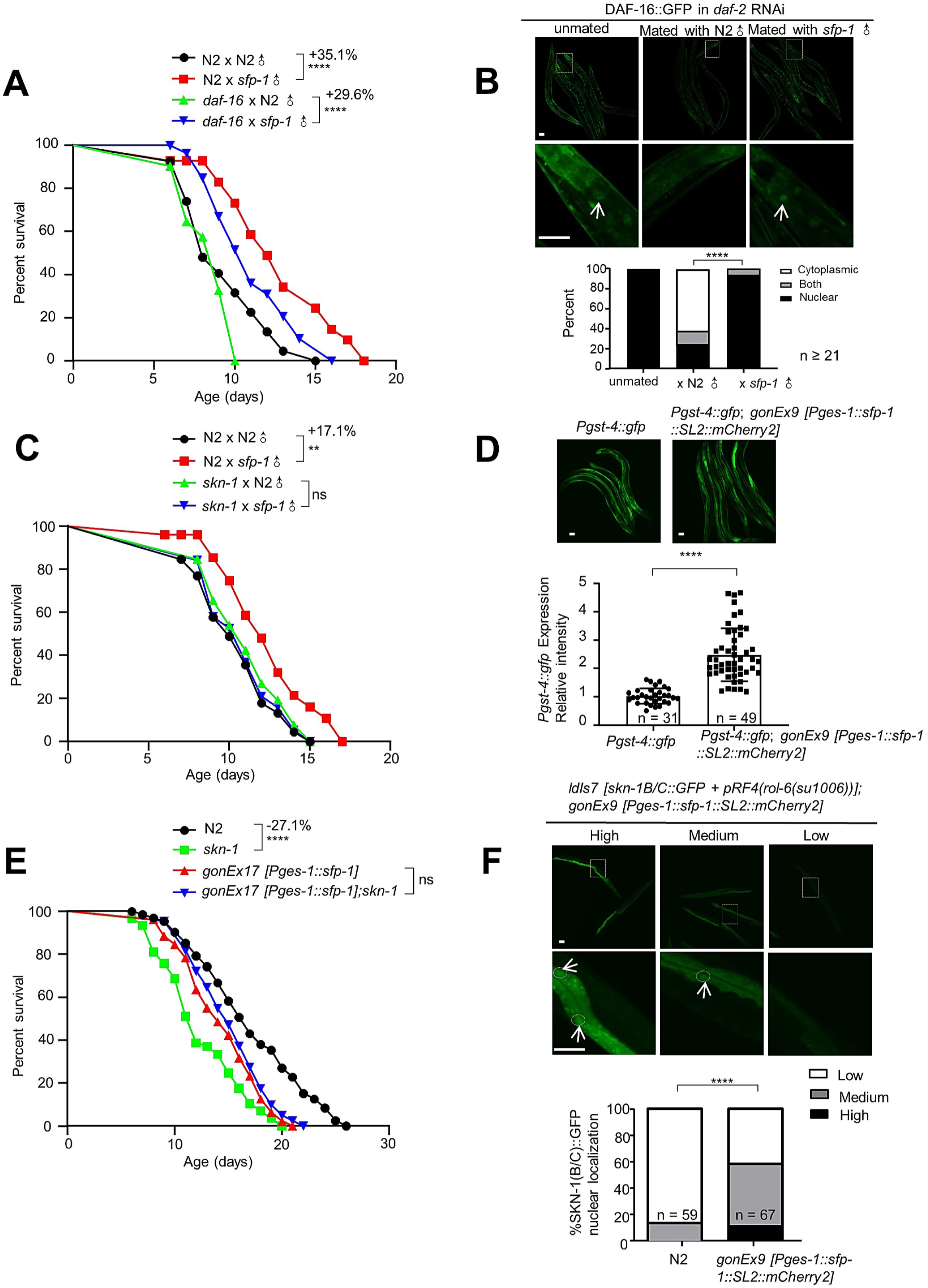
SFP-1 activates the Nrf2 transcription factor homolog SKN-1 after mating. (A) Lifespan survival curves of WT animals and *daf-16* mutant animals treated with WT males or *sfp-1* males. Mean lifespans of unmated and mated N2 and *daf-16*. N2 x N2: 9.33 ± 0.49 days, (n = 25); N2 x *sfp-1*: 12.60 ± 0.72 days, (n = 22); *daf-16* x N2: 8.55 ± 0.36 days, (n = 12); *daf-16* x *sfp-1*: 11.08 ± 0.55 days, (n = 21). \*\*\*\**P* < 0.0001. (B) DAF-16::GFP localization at adult Day 5 and cultured on *daf-2* RNAi bacteria. 10x; Quantitation of DAF-16::GFP, *P* ˂ 0.001. Each worm was assigned a category based on DAF-16::GFP localization. Chi-square test was used to determine the significance. (Scale bars: 100 μm,15 μm) (C) Lifespan survival curves of WT animals and *skn-1* mutant animals treated with WT males or *sfp-1* males. Mean lifespans of unmated and mated N2 and *daf-16*. N2 x N2: 10.39 ± 0.48 days, (n = 24); N2 x *sfp-1*: 12.16 ± 0.69 days, (n = 17); *skn-1* x N2: 11.00 ± 0.45 days, (n = 26); *skn-1* x *sfp-1*: 10.74 ± 0.51 days, (n = 19). (D) Fluorescent images of SKN-1 reporter animals in wild-type and SFP-1 expressed ectopically worms. (Scale bars: 100 μm) (D) Quantification of SKN-1 activation (SFP-1 expressed ectopically worms normalized to mean of wildtype) from (C), Data shown are representative of n = 3 biological replicates with n > 30 animals per condition for each replicate. \*\*\*\**P* < 0.0001 using t-test. (E) The short-lived life span phenotype of *intestine::sfp-1* transgenic worms can be suppressed by *skn-1* mutation. (F) Effect of peptide *intestine::sfp-1* on nuclear localization of SKN-1.Day 1 *ldIs7 [skn-1B/C::GFP+pRF4(rol-6(su1006))];gonEx9[Pges-1::sfp-1::SL2::mCherry2]* nematodes were visualized for subcellular localization of SKN-1::GFP. Top panel: Representative fluorescence micrographs showing “low,” “medium,” and “high” nuclear accumulation of SKN-1::GFP, respectively. White circles show nuclear locations. Bottom panel: Percentage of nematodes with “low,” “medium,” and “high” nuclear SKN-1::GFP accumulation, respectively. Data are from three independent experiments. Chi-square test was used to determine the significance. (Scale bars: 100 μm).

### SFP-1 activates the Nrf2 transcription factor homolog SKN-1 after mating

The DAF-16/FOXO translocation assay results suggested that SFP-1 may be associated with multiple nucleus location factors to mediate post-mating phenotypes. Using the intestinal over-expressed SFP-1 transgenic line, we conducted RNAi screening for transcription factors that were known to be primarily involved in the regulation of longevity and metabolism to identify other important transcription factors (**Figure S4A and S4B**). When comparing the mated *skn-1* loss-function mutant hermaphrodite with the wild-type male, we discovered that the *sfp-1* mutant male did not affect the shorter lifespan phenotype (**Figure 5C**). While overexpressing SFP-1 in the gut cells did not further shorten the lifespan which was the case with the other RNAi transcription factors, however, it slightly extended the lifespan—even the *skn-1* mutant had a shorter lifespan (**Figure 5E**). Moreover, SFP-1 was able to enhance the SKN-1::GFP expression level, including the SKN-1 signal downstream reporter gene *gst-4* (**Figure 5D and 5F**). Sustainably activating the GST protein family contributes to negative effects on longevity ^45^. Thus, even though the over-expressed SFP-1 transgenic line had active *gst-4*, the strain had a shortened lifespan. Meanwhile, the RNAi knock-down of the SKN-1 signal pathway major involving genes such as *sek-1* and *pmk-1* increased the lifespan of over-expressed SFP-1 transgenic line which was similar to the *skn-1* mutant phenotype ^46^ (**Figure S4C and S4D**). We also predicted the interface detail of SFP-1 and SKN-1 binding complexes. We found that many residues formed a variety of interaction bands between SFP-1 and SKN-1 (**Figure S4F**), indicating SFP-1 could directly contact with SKN-1 to act as a regulator. These results suggest that SFP-1 activates the metabolism transcription factor SKN-1 signal pathway to regulate post-mating lifespan via metabolic dependence contribution.

### SFP-1 regulates post-mating lipid metabolism and triggers Asdf through SKN-1

The hermaphrodites lose their body fat after mating, which contributes to the post-mating death. Previous studies showed that mating-induced fat loss was mediated via seminal fluid ^11^. We found that SFP-1 increased the activity of the metabolism transcription factor SKN-1, which indicates that SFP-1 might be the key component in the seminal fluid that promotes mating-induced fat loss. We performed fatty acid staining and TAG analysis in the hermaphrodites mated with wild-type males and *sfp-1* mutant males. The hermaphrodites mated with mutant reduced fat loss compared with those mated with wild-type, in contrast, after mating with *sfp-1::yfp* males which over-expresses the SFP-1 under its own promoter, showed accelerated fat loss. (**Figure 6A and 6B**, **Figure S5A**). In addition, SFP-1 was able to trigger the fat loss phenotype through the intestine cell in the transgenic animal even without mating (**Figure 6C and 6D**, **Figure S5A**), indicating that SFP-1 may play an important role in lipid metabolism after mating. To further confirm our findings of the SKN-1 on mating-induced fat loss, we used *skn-1* mutant mated with wild-type and *sfp-1* male. Indeed, *skn-1* mutant effectively prevents the fat loss no matter mated with wildtype or *sfp-1* male (**Figure 6E and 6F**). Furthermore, this transgenic line in *skn-1* mutant background had more fat than wild-type, similar to the *skn-1* mutant animal, suggesting SFP-1 plays no role in inducing fat loss in the absence of SKN-1 (**Figure 6G and 6H**). Surprisingly, when zooming in to observe the staining images, we found that the fatty acid depleted severely in the soma cells than the germline cells in the over-expressed SFP-1 transgenic line wildtype background whereas the *skn-1* mutant did not, which is known as age-dependent somatic depletion of fat (Asdf) ^47,48^ (**Figure 6I and 6J**). This phenotype is triggered by SKN-1 over-activation or age-related lipid metabolism indeed worms were in the adult day 4 stage not aging yet. Asdf is mediated by vitellogenin family proteins ^47^, we used RNAi to knock down the genes in this family, and the data showed no more shortened lifespan and Asdf phenotype in the over-expressed SFP-1 transgenic line (**Figure S5B-5I**). These results imply that SFP-1 over-turns on SKN-1 activation to contribute to the post-mating lipid metabolism in mated hermaphrodite gut cells.

**Figure 6:**
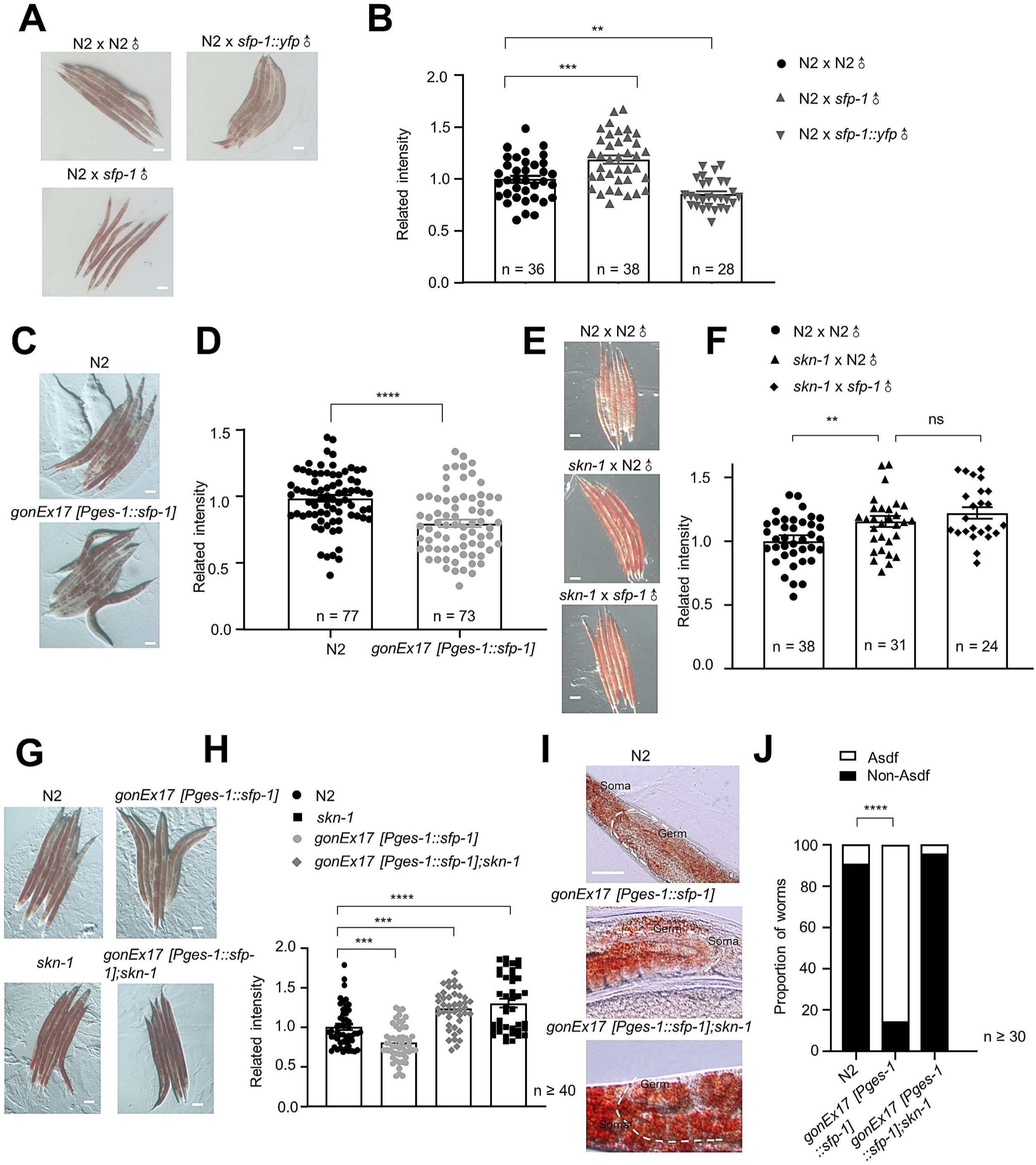
SFP-1 regulates post-mating lipid metabolism and triggers ASDF through SKN-1. (A-B) Representative pictures of Oil Red O staining Day 5 hermaphrodites in the presence of males starting at adult Day 1. Quantification of Oil Red O fat staining. Compared to mating with N2 males, after mating with *sfp-1* males, neutral lipid level of N2 significantly increased, \*\*\**P* ˂ 0.001. In contrast, after mating with sfp-1::yfp males, neutral lipid level of N2 significantly decreased. \*\**P* ˂ 0.01. (Scale bars: 100 μm). (C-D) Representative pictures of Oil Red O staining in 5 days adult control and intestinal overexpressing SFP-1 nematodes. Quantification of Oil Red O fat staining. Compared to control, neutral lipid level of overexpressing SFP-1 in intestine animals significantly decreased. \*\*\*\**P* ˂ 0.0001. (Scale bars: 100 μm). (E-F) Representative pictures of Oil Red O staining Day 5 *skn-1* mutant in the presence of males starting at adult Day 1. Quantification of Oil Red O fat staining. There were no significant differences in fat levels after *skn-1* hermaphrodites mating with *sfp-1* male or N2 male. (Scale bars: 100 μm). (G-H) Representative pictures of Oil Red O staining in 5 days old adult control and intestinal overexpressing SFP-1 in *skn-1* mutant nematodes. Quantification of Oil Red O fat staining. Intestinal overexpressing SFP-1 animals had significantly increased neutral lipid levels in the absence of *skn-1* compared to the WT background. \*\*\*\**P* ˂ 0.0001. (Scale bars: 100 μm). (I-J) In the absence of *skn-1*, intestinal overexpressing SFP-1 animals can partially suppress somatic lipid depletion (Asdf). (Scale bars: 15 μm)

### SFP-1 leads to PUFA depletion in the intestine cells

Similar to the fat loss phenotype in mated hermaphrodites, ectopic expression of SFP-1 in unmated hermaphrodite intestinal cells promoted depletion of lipids. It is necessary to investigate if SFP-1 accelerates fat consumption or inhibits fat storage. Through lipidomic profiling of FFAs, we found that while PUFAs were significantly reduced by the ectopic expression of SFP-1, SFAs, and MUFAs did not change in content amount (**Figure 7A**). This finding suggests that SFP-1 accelerates the depletion of fat, particularly the PUFA class, but not the intake or storage of fat. To test the hypothesis that PUFAs regulate longevity and fat levels in the ectopic expressed SFP-1 worms. We used RNAi to knock down *fat-1* and *fat-2* that encode ω-3 fatty acid desaturases and Δ12-desaturase, respectively required for PUFA biosynthesis ^49^ (**Figure 7B**). The over-expression of SFP-1 in gut cells was no longer associated with fat loss when these desaturase genes were knocked down, while a noticeable decline in the control worms (**Figure 7C and 7D**). Furthermore, knocking down the desaturase gene also prevented the shortened lifespan in the ectopic expressed SFP-1 worms (**Figure 7E and 7F**). Indeed, compared to empty vector controls, the lifespan of the ectopically expressed SFP-1 worms was increased by RNAi knockdown of other fatty acid desaturases, such as *fat-3, fat-4, fat-5* (**Figure S6A-S6C**). In contrast, we found that *fat-6* RNAi significantly reduced the lifespan of SFP-1 expressed ectopically worms (**Figure S6D**). *fat-6* processed the synthesis of MUFAs which are the substrates of fatty acid desaturases, therefore, knocking down the gene had the parallel effect on lifespan regulation. Meanwhile, *fat-7* RNAi had almost no effect on the lifespan of SFP-1 expressed ectopically worms (**Figure S6E**). Together, these results suggest that SFP-1 triggers PUFA depletion in intestine cells, which causes post-mating fat loss in mated animals contributing to the shortened lifespan.

**Figure 7:**
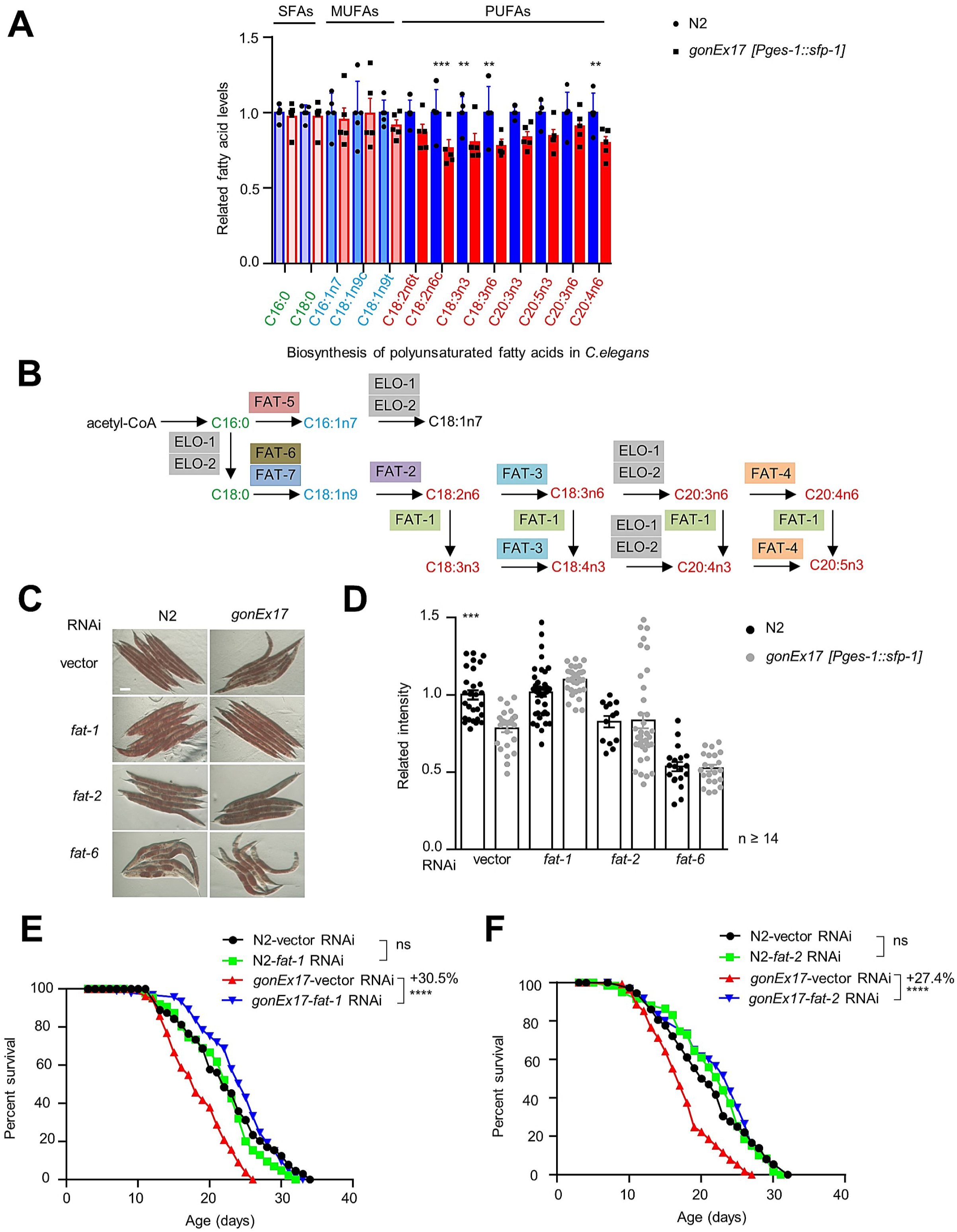
SFP-1 triggers PUFAs depletion in intestine cells. (A) Fatty acid profile by GC-MS following intestinal overexpressing SFP-1. Fatty acid levels were normalized to the control condition. Data are the mean ± SEM of three independent experiments, each with three biological replicates. Significant *P* values are shown. *P* values: two-way ANOVA with Bonferroni’s multiple comparison test. (B) The pathway of PUFA synthesis in *Caenorhabditis elegans*. (C-D) Oil red O staining (C) and quantification (D) in 5 days adult control and intestinal overexpressing SFP-1 nematodes in *fat* genes RNAi treatments. (D) *intestine::sfp-1* transgenic worms stored less fat when grown on EV. This phenotype was abolished in *fat* genes RNAi treatments. Error bars: SEM; n ≥ 14; \*\**P* < 0.001; t-test. (Scale bars: 100 μm). (E-F) Lifespan survival curves of WT animals and intestinal overexpressing SFP-1 animals treated with EV or *fat* RNAi. The short-lived life span phenotype *of intestine::sfp-1* transgenic worms can be suppressed by *fat-1* (E)*, fat-2* (F), gene knockdown.

## Discussion

We carried out this study to investigate the mechanism of *C. elegans* hermaphrodite post-mating physiological changes regulated through a male seminal fluid protein, SFP-1, which functions as an interactive factor. SFP-1 was expressed in the seminal vesicle and collected into the microvesicle exophers. We had directly observed the transfer of SFP-1 from the hermaphrodite’s somatic gonad tissue uterus and spread to the intestinal cells after copulation. This process required the membrane fusion between the vesicles or vesicle to the membrane in the male seminal fluid or the hermaphrodite’s gonad basal membrane. In mated hermaphrodites, SFP-1 was released from the reproductive tract and may have been taken up into the intestinal cells by endocytosis. Together, these data strongly support a model in which SFP-1 is a component of the male seminal fluid that is transferred during copulation and long-distance targeting to hermaphrodite gut cells via exophers releasing, membrane fusion, and endocytosis, ultimately causing the hermaphrodite post-mating physiological changes. Thus, it is demonstrated that SFP-1 contributes to promoting sex interaction antagonistic responses.

Male sperm accentuated the post-reproduction shortened lifespan through shrinking that is dependent on the germline and DAF-9/DAF-12 ^11^. Male pheromones decreased longevity in the opposite sex by hijacking several well-known longevity pathways ^21,26,50^. However, the effective factor of the receipt and storage of male seminal fluid is not well-known. We discovered the male protein SFP-1, the third way to mediate post-mating phenotype, associated with DAF-16 and SKN-1 transcription factors to play key roles in the substantial cost of mated hermaphrodites. Our study completed the mechanisms of male-induced phenotypes that exist in *C. elegans*.

The over-expression of *sfp-1* in intestinal cells had the opposite effect, promoting a shorter lifespan in the transgenic line, while the *sfp-1* mutant male inhibited the mating-induced shortened lifespan of hermaphrodites. We observed SFP-1 located in the cytoplasm and nucleus in the transgenic worms. We predicted that SFP-1 could bind with DAF-16 and SKN-1, which are important transcription factors in metabolism, stress, and longevity regulation ^51–56^. Therefore, SFP-1 could function as a factor to regulate their translocations. Our data suggested that SFP-1 assisted DAF-16 in moving out from the nucleus and could also maintain it in the cytoplasm so that enhanced post-mated shortened lifespan. On the other hand, previous studies showed that activation of endogenous SKN-1 resulted in decreased survival ^45,47,48^. Over-expressing SFP-1 in gut cells turned on SKN-1 expression and promoted its nucleus translocation, by contrast, knocking down SKN-1 could erase the effect of lifespan shortening caused by SFP-1, thus we assumed that SFP-1 and SKN-1 function together to shorten lifespan.

Our data suggested that the main cause of the reduced fatty acid in mated hermaphrodites could be the SKN-1 over activation induced by seminal fluid protein SFP-1. Additionally, ASDF characterization—the typical lipid metabolism aberrant phenotype of the *skn-1 gf* mutation—can be observed in the over-expressed SFP-1 worm. Mated hermaphrodites also exhibit this phenotype, and the remarkable lipid consumption in these animals is suppressed by mating with the *sfp-1* mutant males. That may be the reason why the brood size of hermaphrodites mated with *sfp-1* mutant male was decreased when compared with mated wildtype male, due to less energy and nutrients transporting from soma cells to the germline. Therefore, SFP-1 improves SKN-1 activity to regulate post-mating lipid metabolism.

Surprisingly, over-expressed SFP-1 in wild-type worm intestine cells can induce sperm-germline dependent shrinking and brood size increase, which was similar to the post-mating phenotypes. Functional male sperms are required for the shrinking in mated hermaphrodites, however, this special ability in male sperm is largely unknown. It is possible that over-expressed SFP-1 applied the gut cell metabolites to germline stimulating hermaphrodite self-sperm conferring some male sperm abilities to induce its own shrinking phenotype. Besides, reproduction activity needs support from somatic cells such as the intestine ^57^. However, not much research indicates that gut cells can directly initiate the reproductive process. We observed that PUFAs, one important type of fat acid in many physiological processes ^58,59^, depleted in the somatic tissues in SFP-1 expressed ectopically worms. There is a possibility that over-expressed SFP-1 exploited the PUFAs to initiate and promote reproduction activity for having more gametes. To properly support the downstream reproduction tissue, SFP-1 triggered the somatic tissue, which acts as the upstream energy station by using fatty acids. That is why the mutants without the germline also lose fat in response to mating ^11^. SFP-1 is the potential programmer of fat depletion to anticipate future resource allocation after mating.

## Materials and Methods

### *C. elegans* Strains and Maintenance

*C. elegans* strains (Table 1) were nurtured at a temperature of 20 °C on plates containing a nematode growth medium (NGM) with OP50 bacteria as the seeding source. The creation of transgenic lines involved the direct injection of plasmid DNA into the gonads of hermaphrodites. To generate SFP-1 fusion-expressing nematodes, 5kb upstream of the *sfp-1* gene was selected as a promoter for expression of SFP-1, fused with YFP, and the coding sequences of *sfp-1* promoter and *sfp-1* tagged with YFP at the C-terminus were cloned into the vector PBS77.To generate tissue-specific SFP-1 fusion-expressing nematodes, *Pges-1*, and *Pmyo-3* as tissue-specific promoter for expression of SFP-1, fused with YFP, and the coding sequences of sfp-1 promoter and sfp-1 tagged with YFP at the C-terminus were cloned into the vector PBS77.To generate tissue-specific expression of SFP-1, the coding sequence of *sfp-1* was cloned between the tissue-specific promoter and the SL2::YFP marker and used for DNA microinjection (50ng/μL) to generate nematodes carrying the extra-chromosomal transgene array.

### CRISPR–Cas9

*sfp-1(syb1800)* was generated by SunyBiotech upon our request. sgRNA was designed according to the required mutant genes, and cas9 sgRNA expression plasmids were constructed. The constructed plasmids were injected into the gonad using a microinjection. PCR and sequencing were used to determine if the gene was successfully recombined and if it had mutated.

### Lifespan assay

Lifespan assays were performed on 60 mm NGM plates at 20℃. ∼100 worms of each genotype were used for lifespan assays and transferred every other day to fresh NGM plates. Survival rate was scored every 1–2 days. Worms were censored if they crawled off the plate, bagged, or exhibited protruding vulva. In all cases, the first day of adulthood was scored as day 1. Lifespan data was analyzed with GraphPad Prism 8 (GraphPad Software, Inc.) and IBM SPSS Statistics 21 (IBM, Inc.). Log-rank (Kaplan-Meier) was used to calculate p values. All lifespan experiments were repeated at least twice. Detailed data analysis is in Supplemental Table S1.

### RNAi treatment

Freshly streaked single colonies of HT115 bacteria containing either empty vector L4440 or RNAi plasmid were grown up overnight (20–24 hr) at 37°C in LB supplemented with carbenicillin (100 mg/ml). The next day, bacteria were diluted into fresh LB and cultured for a few hours, and subsequently, IPTG (200 mM final concentration) was added in liquid culture when bacterial density reached OD600 = 0.6 to induce the production of dsRNA for 4 hr at 37°C. Two days before experiments, freshly grown RNAi bacteria were seeded on RNAi nematode growth media (NGM) plates supplemented with carbenicillin (25 mg/ml) and IPTG (1 mM). All RNAi feeding assays were started from eggs throughout lifespan at 20°C. Tissue-specific gene knockdown using the strains targeting RNAi to the intestine (VP303), Synchronized eggs of VP303 were cultured on NGM plates seeded with E. coli HT115 RNAi clones at 20°C until the progeny reached the L4 stage.

### Mating assay

3.5-cm NGM plate was seeded with 20ul OP50. One synchronized day 2 adult hermaphrodite and three young males (day 1 - day 2) were transferred into each well of the plates and mating was allowed for 24 hours. On day 3, Each hermaphrodite was numbered for experiment and was transferred onto newly NGM plates every day. The old plates were kept for 3 days to check the male/hermaphrodite ratio. The unsuccessfully mated worms were censored from the experiments.

### Brood assay

The nematodes were distributed onto NGM plates with a diameter of 30 mm and one nematode per plate. A minimum of 10 plates were utilized for each brood assay experiment. Egg production and hatching rates were quantified daily, and the entire experiment was replicated three times. *P* values were calculated by one-way ANOVA, Bonferroni’s multiple comparisons test.

### Body length measurement

The middle line of each worm was delineated using the segmented line tool and the total length was documented as the body length of the worm. T-test was performed to compare the body size difference between groups of worms on the same day.

### Confocal microscopy

Live worms were mounted on 2% agarose pads with 20 mM levamisole and examined using an Olympus FV3000 scanning confocal microscope.

For imaging of SFP-1 release during mating, three *glo-4* hermaphrodites were placed with ten males of the *sfp-1::yfp* strain in 3.5-cm NGM with 20ul OP50. The mating process lasts for 30 minutes. The mated hermaphrodites were imaged 0.1h and 0.5h after mating. Imaging was performed using 60x objective.

### Imaging analysis

To quantify *sfp-1::yfp* fluorescence intensity, 20-30 worms of each group were anesthetized with 20 mM levamisole on 2% agarose pads. Images were acquired on FV3000 (Olympus) under 10X objective and analyzed with ImageJ (NIH). Image J was used to measure the mean and the maximum GFP intensity of either the seminal vesicle area. T-test analysis was performed to compare the GFP intensity of different group of worms.To quantify *gst-4*::*gfp* fluorescence intensity, at least 30 worms day 5 adult worms were anesthetized with 20 mM levamisole on 2% agarose pads. Images were acquired on FV3000 (Olympus) under 10x objective and analyzed with ImageJ (NIH). T-test analysis was performed to compare the GFP intensity of different group of worms.

### Scoring exophers and fluorescence microscopy

The representative pictures presented in the manuscript were acquired with an inverted FV3000 confocal microscope with a 60x oil immersion objective. For each exopher scoring assay, on adulthood day 2, animals were visualized on NGM plates, and the number of exophers was counted in each freely moving worm. *P* values were calculated by one-way ANOVA, Bonferroni’s multiple comparisons test.

### Oil Red O staining and quantification

Mid-L4 stage animals were placed on OP50 NGM plates and raised to day 5 adult stage. Worms were washed off plates with PBS+triton, then rocked for 3 min in 60% isopropyl alcohol before being pelleted and treated with ORO in diH2O overnight.

Worms were washed in PBS+triton three times before being imaged at 5× magnification with Mshot MS23 software and ZEISS steREO Discovery V8 color camera. For lipid content quantification, color images were background subtracted and converted into grayscale images using ImageJ program. Signal intensities were then compared among worms grown under different conditions. At least 30 worms were assayed for each experiment. T-test analysis was performed to compare the fat staining of two groups of worms. When more than two groups of worms, *P* values were calculated by one-way ANOVA, Bonferroni’s multiple comparisons test.

### DAPI staining and counting of intestine nucleus

Worms were stained according to the Bio-protocol (http://www.bioprotocol.org/wenzhang.aspx?id=77). At least 20 worms day 1 adult worms were anesthetized with 20 mM levamisole on 2% agarose pads. Images were taken with an FV3000. Images were acquired on FV3000 (Olympus) under 60x objective. Intestinal cell nucleus are identified by DAPI staining. Photos are taken and the number of intestinal nucleus within the merge image is counted. T-test analysis was performed to compare the GFP intensity of different group of worms. *P* values were obtained by T-test.

### MitoTracker Staining

MitoTracker Red CMXRos (Invitrogen) dye, which selectively stains mitochondria, was used for labeling *sfp-1::yfp* male sperm, MitoTracker does not affect sperm function or motility. 100–150 males were transferred to a watchglass containing 10 μM MitoTracker in 300 μl M9 buffer. Males were incubated in the dark for 4 hours, transferred to an NGM plate mated with *glo-4*.

### SKN-1 Nuclear localization assay

SKN-1 nuclear localization was analyzed using the transgenic LG326 nematode that expresses a fusion protein of SKN-1 tagged with a GFP reporter. Fluorescent images were captured by using FV3000 (Olympus) with 20x magnification. For quantification, the LG326 nematodes were categorized into three groups (i.e., “low,” “medium,” and “high”) depending on the levels of SKN-1::GFP nuclear accumulation in the intestinal nuclei. “Low” represents that SKN-1::GFP nuclear accumulation is barely detectable through the intestine, “medium” refers to the nematodes in which SKN-1::GFP nuclear ac-cumulation is partially present in intestine, and “high” indicates that a strong SKN-1::GFP signal is observed in almost all intestinal nuclei ^56^. Day 1 adult worms (n > 50) per strain in each experiment were randomly selected to score sub-cellular localization of GFP-fused proteins, and the percentage of nematodes in each category was calculated. All experiments were repeated independently at least two times with similar results. Chi-square test was used to determine the significance.

### FOXO/DAF-16 localization assay

Hermaphrodites were synchronized using a timed egg lay on *daf-2* RNAi plates seeded with HT115. coli containing EV. Starting at adult day 1, synchronized hermaphrodites were selected to be either cultured only with other hermaphrodites (around 35 per 6-cm plate) or with males (around 15 hermaphrodites and around 45 adult day 1 *sfp-1* males or N2 males). After 4 days (adult day 5), the worms were immediately anesthetized in M9 containing 20 mM levamisole, mounted on 2% agar pads and imaged. Fluorescent images were captured by using FV3000 (Olympus) with 10x magnification. Individual worms scored as primarily cytoplasmic, primarily nuclear or nuclear and cytoplasmic DAF-16::GFP localization in the anterior intestinal cells. Chi-square test was used to determine the significance.

### Asdf quantification

Worms stained with ORO were photographed using Olympus FV3000 and a 60x oil immersion objective. Worms were scored as previously described ^46^. Fat levels of worms were placed into three categories: non-Asdf, intermediate, and Asdf. Non-Asdf worms display no loss of fat and are stained dark red throughout most of the body (somatic and germ cells). Intermediate worms display significant fat loss from the somatic tissues, with portions of the intestine being clear, but ORO-stained fat deposits are still visible (somatic < germ cells). Asdf worms had most, if not all, observable somatic fat deposits depleted (germ cells only) or significant fat loss from the somatic tissues with portions of the intestine being clear (somatic < germ) ^47^. Chi-square test was used to determine the significance.

### Protein Prediction

Alphafold2 was used to predict the structure of SFP-1, SFP-1 structure prediction file O01420. Model confidence around the caspase cleavage site is very high (pLDDT > 90).

### Protein-protein Interaction Prediction

HDOCK is used for fast protein-protein docking (http://hdock.phys.hust.edu.cn/). The docking program first samples the putative binding modes between two proteins based global search method and then evaluates the sampled modes with a knowledge-based scoring function for protein-protein interactions. The model of SKN-1 or DAF-16 is used as the docking receptor, SFP-1 is used as the ligand for docking. The binding mode of the heterodimer was determined based on the combination of complex structure and scoring function. Visualization and analysis of model features were carried out by Open-Source Pymol (https://pymol.org).

### TAG measurement

Day 5 adult hermaphrodites (n = 100) were collected, washed with M9, and homogenized in 150 uL lysis Buffer. TAG was measured by using a colorimetric assay kit (E1013-50, Applygen). TAG concentrations were normalized relative to protein concentration using the BCA protein assay. *P* values were obtained by T-test.

### Drug treatment assays

For drug treatment, worms were cultured in OP50 with either 0.2% DMSO or 10 μM bafilomycin A1 (dilute with DMSO). day 2 unmated worms were treated on drugged plates for two days, day 2 worms were mated in drugged plates for two days and then transferred to normal NGM and observed for longevity.

### GC–MS analysis of fatty acid profiles

300 age-synchronized old adult (adult day 5) animals were collected in M9 buffer and washed three times to remove residual bacteria in the worm pellets. A small steel ball was added into the sample tube, then homogenating for 30 min. After that, the sample was mixed with 150 μL MeOH, 200 μL MTBE and 50 μL 36% phosphoric acid/water (precooled at −20°C). The suspension was vortexed for 3 min under the condition of 2500 r/min and centrifuged at 12000 r/min for 5 min at 4°C. Take 200 μL of supernatant into a new centrifuge tube blow dry and add 300 μL methanol solution of 15% boron trifluoride, vortex for 3 min under the condition of 2500 r/min, kept in the oven at 60°C for 30 min. Cool to room temperature, then accurately add 500 μL n-hexane and 200 μL saturated sodium chloride solution. After vortexing for 3 min and centrifugation at 4°C and 12000 r/min for 5 min, transfer 100 μL n-hexane layer solution for further GC-MS analysis. The fatty acid concentration of the interventions was normalized to the fatty acid concentration of the control. The final ratio is expressed as relative fatty acid levels in the graph. Each experiment was carried out at least three times independently. Relative fatty acid abundances were plotted using Prism 8 and statistically significant differences between samples were assessed using a two-way analysis of variance with Bonferroni’s multiple comparison test. When the p-value was less than 0.05, there was a significant difference, which was recorded as *. p < 0.01 was marked as **, p < 0.001 was marked as ***, and p < 0.0001 was marked as ****.

## Acknowledgments

We thank Y. Wang, W. Yang, and L. Zhu for technical assistance. We thank Wei Zou from Zhejiang University, Xiajing Tong from Shanghai Science and Technology University, and Shawn Xu from University of Michigan for the assistance and valuable suggestions provided in this study. Some strains were obtained from the Caenorhabditis Genetics Center. This work was funded by the National Natural Science Foundation of China (32100604) and the program for HUST Academic Frontier Youth Team (5001170068).

## Competing interests

The authors report no competing interests.

## Data availability

The datasets generated and analyzed during the current study are available from the corresponding author upon reasonable request.

## Authors’ Contributions

M.C. and J.G. designed the experiments and processed the data. M.C. prepared the figures. M.C. and J.G. analyzed and interpreted the results. J.G. wrote the paper and all authors reviewed and revised the paper. All authors have read and agreed to the published version of the manuscript.

---

**Supplementary Figure 1:**
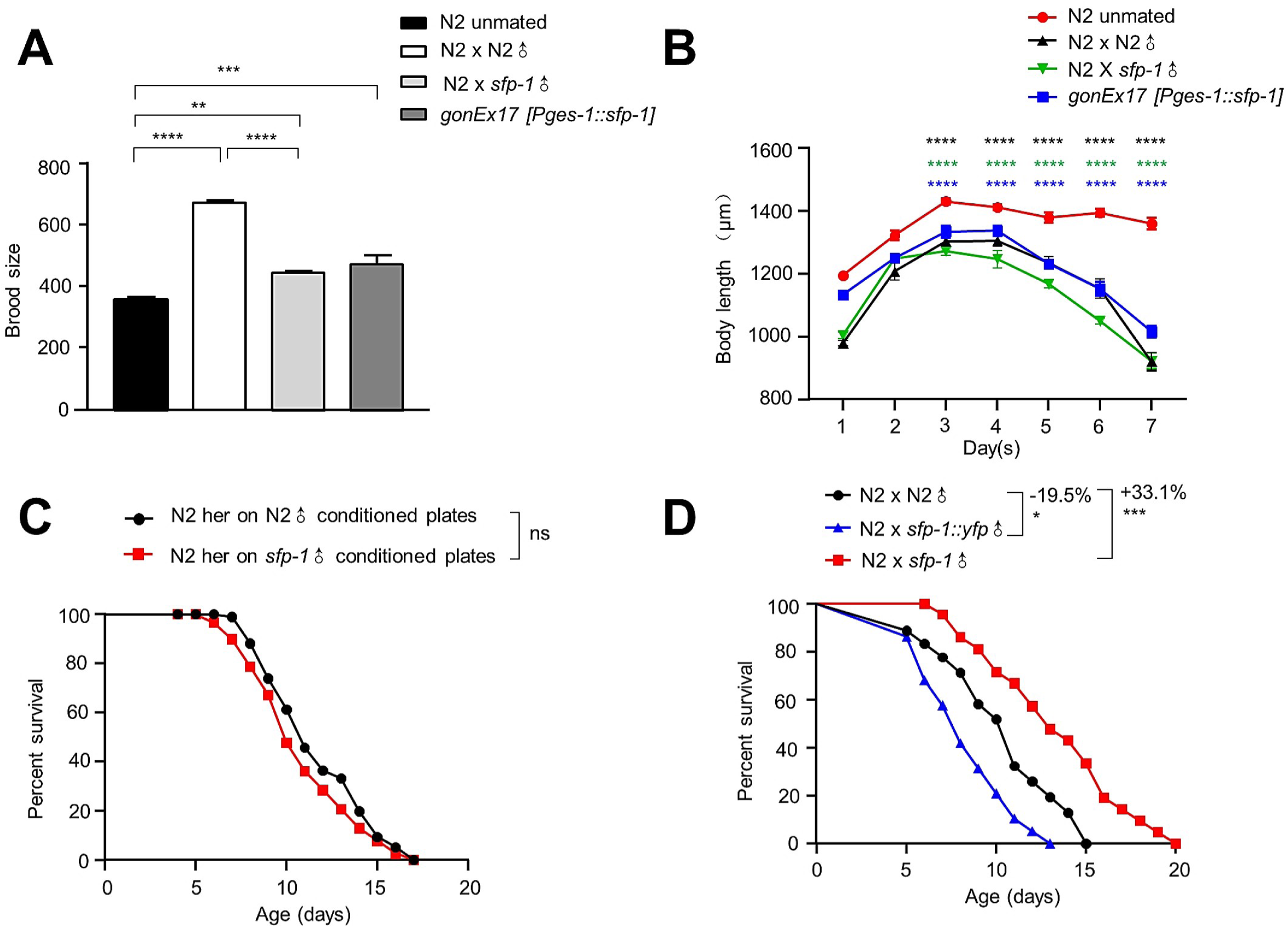
The secreted protein SFP-1 is involved in mating-induced phenotype, *sfp-1* mutant male sperm and pheromones still influence hermaphrodites. (A) Total brood size of N2 worms in different conditions. Fertility was assayed by measuring total offspring production by individual hermaphrodites at 20°C. \*\**P* < 0.01, \*\*\**P* < 0.001, \*\*\*\**P* < 0.0001, n > 15. *P* values were calculated by one-way ANOVA, Bonferroni’s multiple comparisons test. (B) Length of N2 worms in different conditions. \*\*\*\**P* < 0.0001, n > 15. *P* values were calculated by t-test. (C) Lifespan of N2 in N2 male condition plates and *sfp-1* male condition plates. N2 in N2 male condition: 11.72 ± 0.28 days, n = 97 worms; N2 in *sfp-1* male condition: 10.89 ± 0.31 days, n = 79 worms, *P* = 0.066. (D) Lifespan of mated N2 worms. N2 x N2 ♂: 10.22 ± 0.78 days, n = 16 worms; N2 x *sfp-1::yfp* ♂: 8.22 ± 0.54 days, n = 20 worms; N2 x *sfp-1*♂: 12.93 ± 0.93 days, n = 22 worms. \**P* < 0.05, \*\*\**P* ˂ 0.001.

**Supplementary Figure 2:**
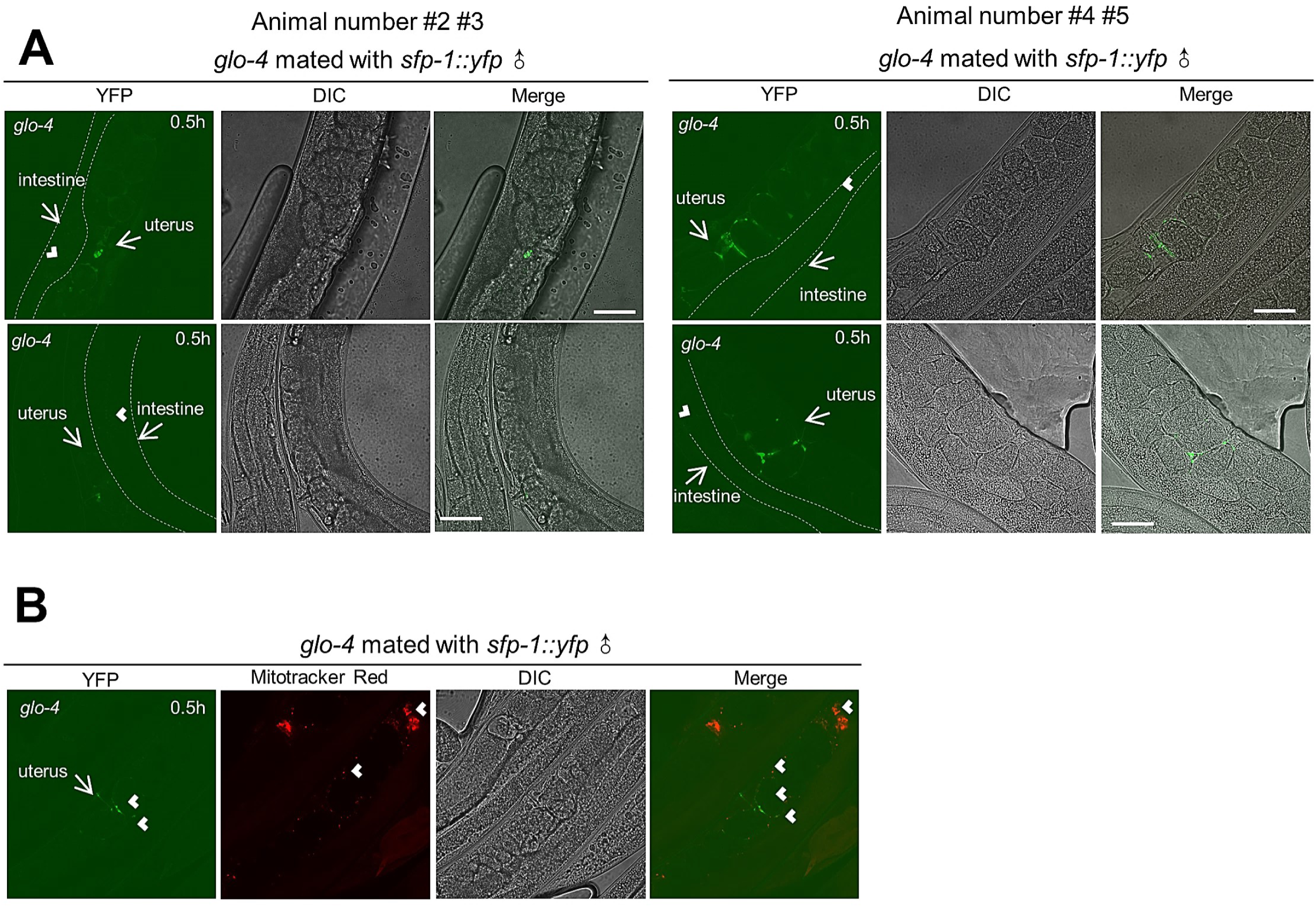
SFP-1 spreads into the intestine cells from the uterus in mated hermaphrodites via endocytosis. (A) Other animals. Male-to-hermaphrodite transfer experiment where *sfp-1::yfp* males were mated with low autofluorescence *glo-4* hermaphrodites. Imaging of the low autofluorescence *glo-4* hermaphrodite after copulation revealed the presence of male derived *sfp-1::yfp* in the uterus, and then half an hour after mating, lots of weak fluorescence appeared in the intestine cell. (B) Male releases *sfp-1::yfp* during sperm transfer. White arrow indicate *sfp-1::yfp* fluorescent signal (green) and sperm (red).

**Supplementary Figure 3:**
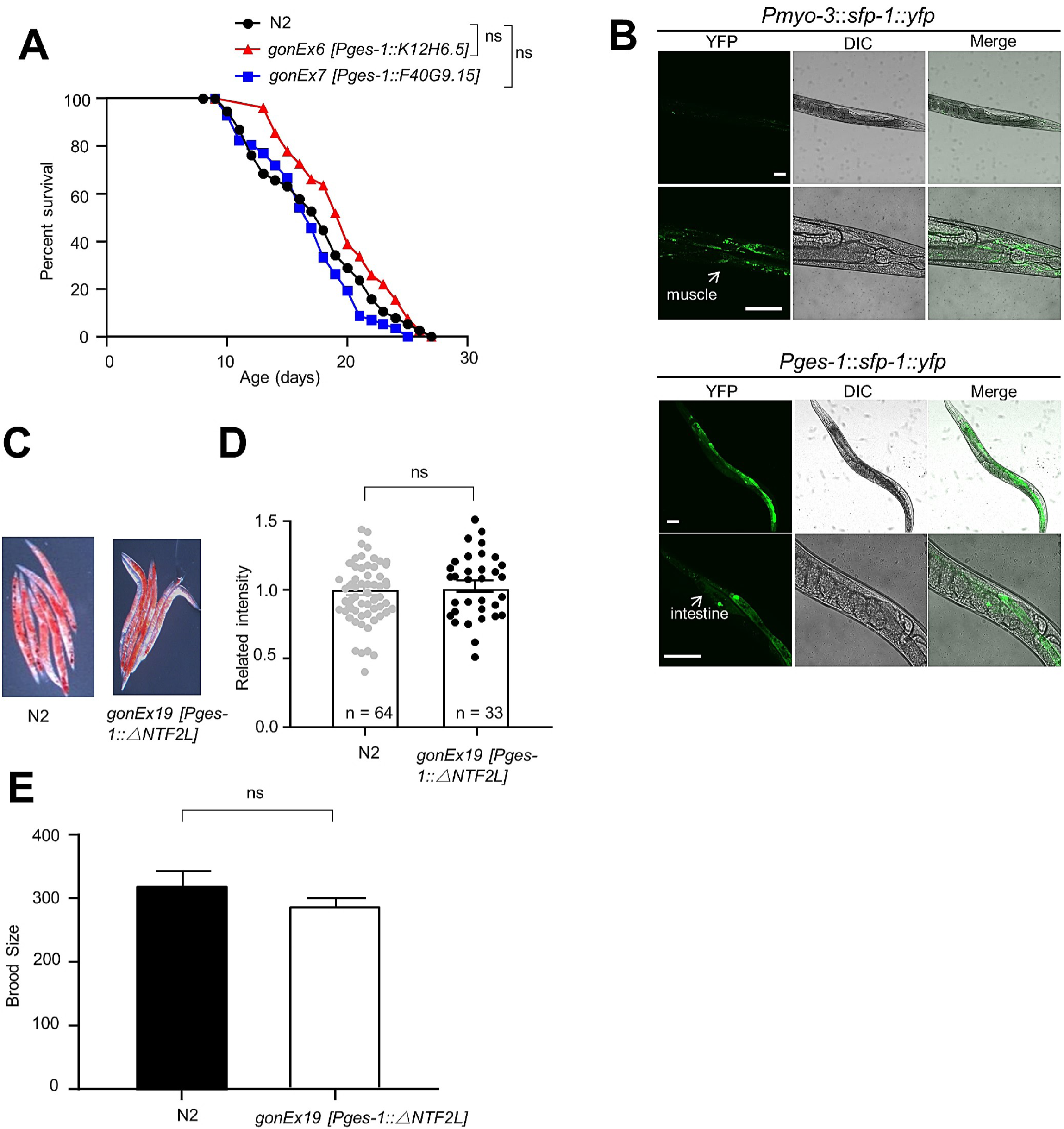
The potential function of NTF2L in longevity, lipid metabolism, and reproduction. (A) Lifespan survival curves of WT animals and intestinal overexpressing K12H6.5 or F40G9.15 animals. The other seminal fluid proteins such as K12H6.5 and F40G9.15 in the intestine had almost no effect on longevity. (B) Expression of *Pmyo-3::sfp-1::yfp* and *Pges-1::sfp-1::yfp*. (Scale bars: 50 μm, 15 μm) (C-D) Representative pictures of Oil Red O staining in 5 days adult control and *gonEx19 [Pges-1::△NTF2]* worms. Quantification of Oil Red O fat staining. Compared to the control, the neutral lipid level difference was abolished in the absence of NTFL. (E) Total brood size of N2 worms and *gonEx19 [Pges-1::△NTF2]* worms. Rather than promoting fertility, as observed for mated N2 and *intestine::sfp-1* transgenic worms, lack of NTF2L markedly inhibits fertility.

**Supplementary Figure 4:**
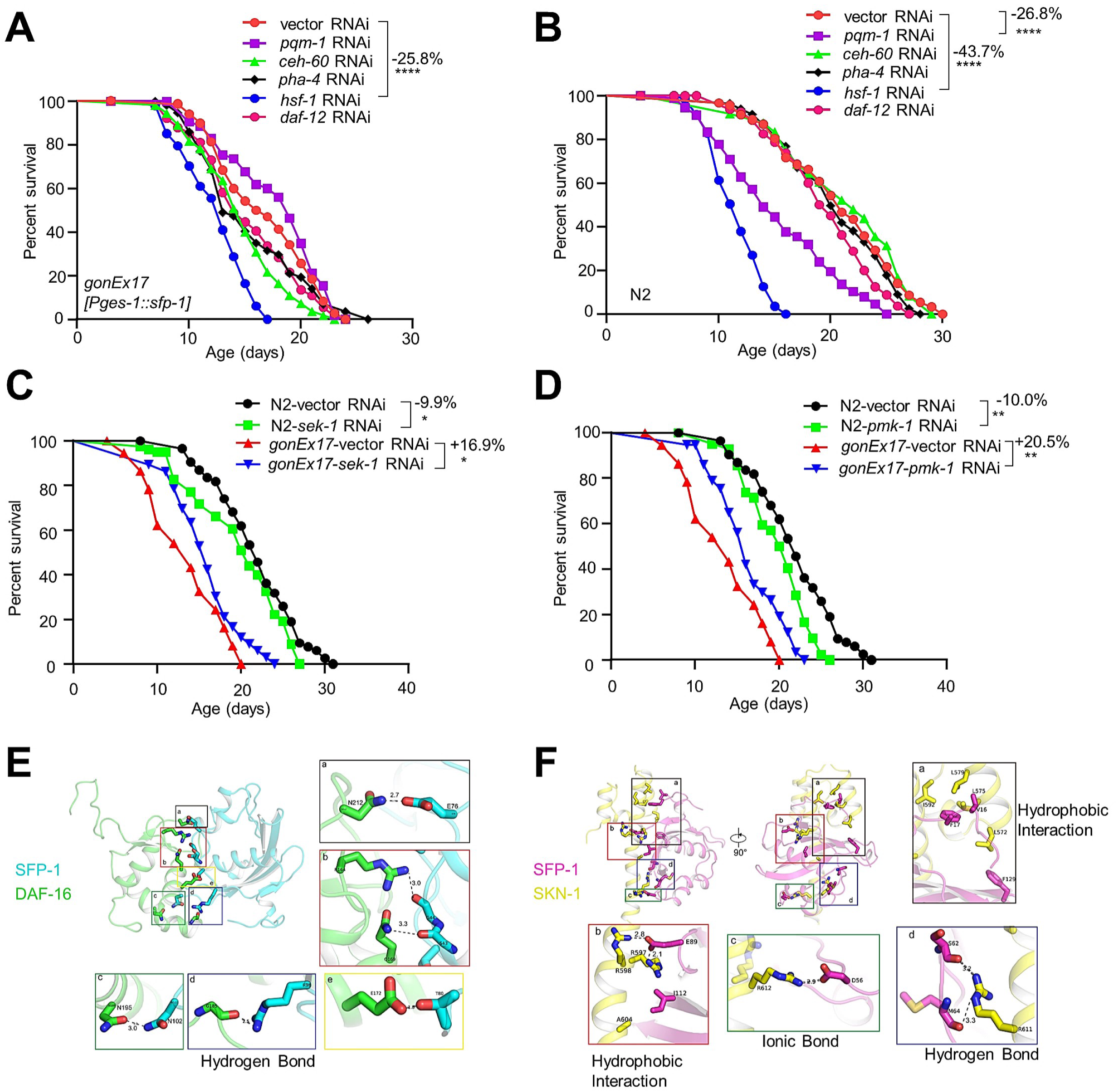
Knockdown of the SKN-1 signal pathway major involving genes increased the lifespan of intestinal overexpressing SFP-1 transgenic line. (A) Lifespan survival curves of intestinal overexpressing SFP-1 animals treated with classical transcription factor genes RNAi. (B) Lifespan survival curves of WT animals treated with classical transcription factor genes RNAi. (C-D) Knockdown of SKN-1 upstream effector *sek-1* (C) and *pmk-1* (D) by RNAi extends the lifespan of SFP-1 expressed ectopically worms. (E) The interaction of SFP-1 with DAF-16. Differently colored proteins denoted different proteins. SFP-1 in blue, DAF-16 in green. Squares show the zoomed interface of the proteins and the hydron bonds between interacting amino acids. (F) The interaction of SFP-1 with SKN-1. Differently colored proteins denoted different proteins. SFP-1 in pink, SKN-1 in yellow. Squares show the zoomed interface of the proteins and the hydron bonds, ionic bonds, and hydrophobic interaction, between interacting amino acids.

**Supplementary Figure 5:**
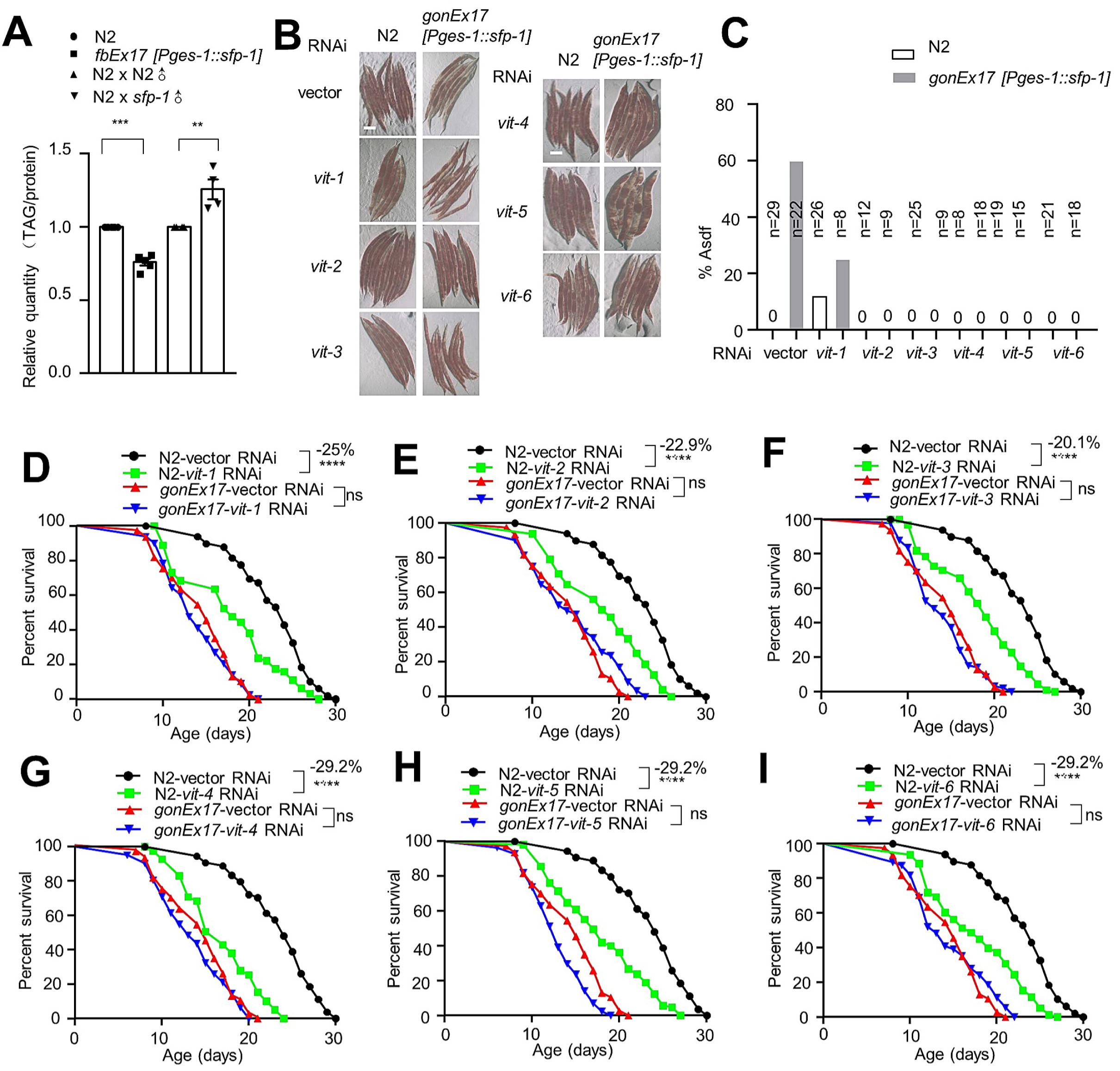
*vit* genes are not involved in the lifespan regulation of intestinal overexpressing SFP-1 animals. (A) Triglyceride levels in N2 worms in different conditions. Data represent mean ± SEM, three biological replicates per group per independent repeat, and three independent experiments. \*\**P* < 0.01, \*\*\**P* < 0.001 and *P* values were obtained by t-test. (B-C) % Asdf is quantified by counting the number of animals treated with EV or *vit* RNAi in a cohort that displays the phenotype compared to the number of animals that do not. (D-I) Lifespan survival curves of WT animals and intestinal overexpressing SFP-1 animals treated with EV or *vit* RNAi. *vit* RNAi had almost no effect on the lifespan of SFP-1 expressed ectopically worms.

**Supplementary Figure 6:**
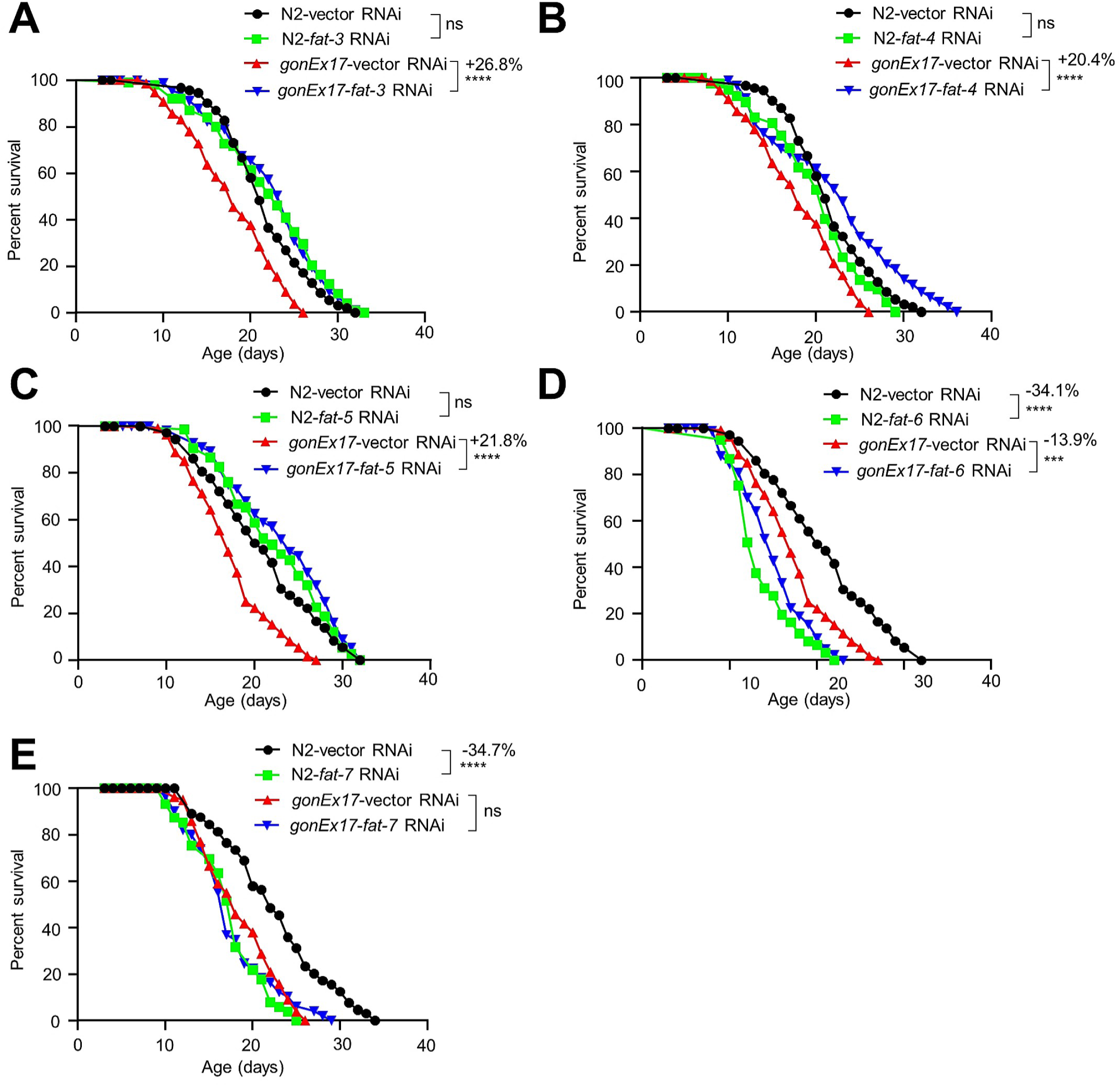
Fatty acid desaturases are involved in the lifespan regulation of intestinal overexpressing SFP-1 animals. (A-C) Lifespan survival curves of WT animals and intestinal overexpressing SFP-1 animals treated with EV or *fat* RNAi. The short-lived life span phenotype of *intestine::sfp-1* transgenic worms can be suppressed by *fat-3* (A)*, fat-4* (B)*, fat-5* (C) gene knockdown. (D-E) Lifespan survival curves of WT animals and intestinal overexpressing SFP-1 animals treated with EV or *fat* RNAi. *fat-6* (D) and *fat-7* (E) RNAi had almost no effect on the lifespan of SFP-1 expressed ectopically worms.

**Supplementary Figure 7:**
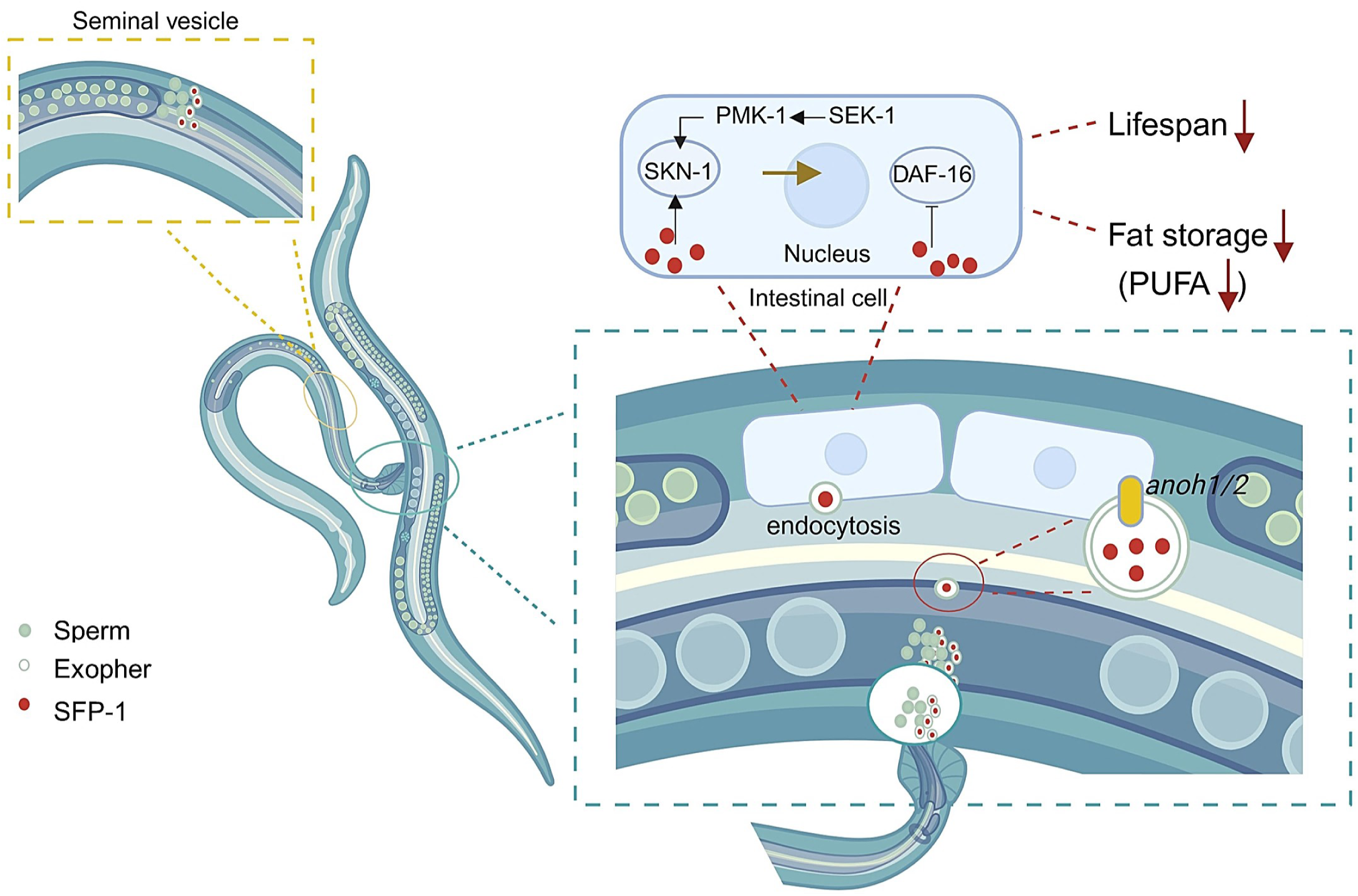
Working model. A schematic model illustrating the trail of SFP-1 transport and the regulation pathways in post-mating longevity and fat metabolism.

## Notes

### Competing Interest Statement

The authors have declared no competing interest.

