## Supplemental Table1 for "A male seminal fluid protein SFP-1 regulates hermaphrodite post-mating longevity and fat metabolism in *Caenorhabditis elegans*"

| **Strain** | **Mean Lifespan ±SEM** | **Median Lifespan** | **75% Lifespan** | **n** | | ***P* value against control** | **Figure** | **Date** |
| --- | --- | --- | --- | --- | --- | --- | --- | --- |
| N2 x N2♂ | 11.01±0.66 | 10 | 8 | 30/40 | | - | 1G | 04-13-2021 |
| N2 x *sfp-1*♂ | 14.98±0.72 | 16 | 11 | 30/39 | | <0.001 | 1G |  |
| N2 x N2♂ | 11.54±0.47 | 11 | 10 | 25/29 | | - | exp. 2 | 10-24-2019 |
| N2 x *sfp-1*♂ | 13.18±0.37 | 13 | 12 | 46/60 | | 0.009 |  |  |
| N2 x N2♂ | 11.48±0.67 | 11 | 10 | 19/27 | | - | exp. 3 | 11-23-2019 |
| N2 x *sfp-1*♂ | 13.58±0.57 | 13 | 13 | 26/30 | | 0.005 |  |  |
| N2 her on N2♂ conditioned plates | 11.72±0.28 | 11 | 9 | 97/118 | | - | S1D | 01-01-2024 |
| N2 her on *sfp-1*♂ conditioned plates | 10.89±0.31 | 10 | 9 | 79/96 | | 0.066 | S1D |  |
| N2 her on N2♂ conditioned plates | 11.62±0.66 | 12 | 9 | 42/42 | | - | exp. 2 | 01-06-2024 |
| N2 her on *sfp-1*♂ conditioned plates | 13.01±0.75 | 13 | 10 | 28/33 | | 0.197 |  |  |
| N2 x N2♂ | 10.22±0.78 | 12 | 10 | 16/18 | | - | S1F | 05-03-2024 |
| N2 x *sfp-1::yfp*♂ | 8.22±0.54 | 8 | 6 | 20/22 | | 0.025 | S1F |  |
| N2 x *sfp-1*♂ | 13.60±0.80 | 13 | 10 | 20/22 | | 0.003 | S1F |  |
| N2 x N2♂  N2 x *sfp-1::yfp*♂  N2 x *sfp-1*♂ | 11.20±0.56  9.49±0.42  12.93±0.93 | 11  9  12 | 9  8  10 | 21/24  34/40  22/27 | | -  0.013  0.005 | exp. 2 | 05-03-2024 |
| control- unmated | 15.63±0.97 | 15 | 12 | 19/26 | | - | 2B | 04-03-2024 |
| bafA1- unmated | 12.65±0.79 | 12 | 10 | 20/29 | | 0.015 | 2B |  |
| control- mated | 7.72±0.35 | 7 | 7 | 20/28 | | <0.0001 | 2B |  |
| bafA1- mated | 9.68±0.44 | 10 | 8 | 21/30 | | <0.0001 | 2B |  |
| control- unmated | 16.38±1.41 | 14 | 11 | 16/19 | | - | exp. 2 | 04-28-2024 |
| bafA1- unmated | 13.72±0.86 | 13 | 10 | 20/23 | | 0.049 |  |  |
| control- mated | 10.02±0.59 | 10 | 8 | 13/19 | | <0.0001 |  |  |
| bafA1- mated | 12.81±0.63 | 12 | 10 | 16/19 | | 0.019 |  |  |
| control- unmated | 16.95±0.95 | 16 | 13 | 23/26 | | - | exp. 3 | 05-01-2024 |
| bafA1- unmated | 14.11±0.65 | 14 | 11 | 18/20 | | 0.003 |  |  |
| control- mated | 11.16±0.49 | 11 | 10 | 22/24 | | <0.0001 |  |  |
| bafA1- mated | 14.09±0.58 | 13 | 10 | 18/23 | | <0.0001 |  |  |
| VP303-vector RNAi - unmated | 17.40±0.31 | 18 | 14 | 20/24 | | - | 2C | 05-23-2024 |
| VP303-*rab-10* RNAi -unmated | 15.84±0.65 | 16 | 12 | 44/50 | | 0.287 | 2C |  |
| VP303-vector RNAi - mated | 9.63±0.85 | 10 | 7 | 16/25 | | <0.0001 | 2C |  |
| VP303-*rab-10* RNAi -mated | 13.49±0.51 | 13 | 11 | 39/45 | | 0.006 | 2C |  |
| VP303-vector RNAi -unmated | 19.00±0.86 | 18 | 16 | 35/44 | | - | exp. 2 | 06-03-2024 |
| VP303-*rab-10* RNAi -unmated | 17.52±1.17 | 17 | 14 | 17/19 | | 0.067 |  |  |
| VP303-vector RNAi - mated | 11.16±0.38 | 10 | 8 | 57/65 | | <0.0001 |  |  |
| VP303-*rab-10* RNAi -mated | 12.89±0.66 | 12 | 9 | 39/45 | | <0.0001 |  |  |
| VP303-vector RNAi - unmated | 17.40±0.31 | 18 | 14 | 20/24 | | - | 2D | 05-23-2024 |
| VP303-*cav-1* RNAi -unmated | 17.76±1.06 | 18 | 15 | 17/24 | | 0.941 | 2D |  |
| VP303-vector RNAi - mated | 9.63±0.85 | 10 | 7 | 16/25 | | <0.0001 | 2D |  |
| VP303-*cav-1* RNAi -mated | 12.76±0.62 | 13 | 9 | 50/65 | | 0.004 | 2D |  |
| N2 x N2♂ | 10.45±0.64 | 10 | 8 | 22/29 | | - | 3C | 09-03-2023 |
| N2 X *anoh-1*♂ | 10.52±0.56 | 10 | 8 | 23/39 | | 0.90 | 3C |  |
| N2 X *anoh-2*♂ | 11.93±0.93 | 10 | 9 | 20/36 | | 0.09 | 3C |  |
| N2 X *anoh-1;anoh-2*♂ | 16.26±0.91 | 16 | 13 | 22/34 | | <0.0001 | 3C |  |
| N2 x N2♂ | 9.88±0.47 | 9 | 8 | 24/29 | | - | exp. 2 | 09-15-2023 |
| N2 X *anoh-1*♂ | 11.84±0.59 | 11 | 10 | 16/28 | | 0.035 |  |  |
| N2 X *anoh-2*♂ | 11.43±0.78 | 10 | 9 | 30/40 | | 0.073 |  |  |
| N2 X *anoh-1;anoh-2*♂ | 13.40±0.99 | 14 | 11 | 22/39 | | 0.001 |  |  |
| N2 | 17.90±0.57 | 18 | 14 | 82/102 | | - | 4A | 04-05-2022 |
| *gonEx17 [Pges-1::sfp-1]* | 13.77±0.41 | 13 | 11 | 85/92 | | <0.0001 | 4A |  |
| *gonEx18 [Psmu-1::sfp-1]* | 17.44±0.61 | 17 | 12 | 80/97 | | 0.529 | 4A |  |
| *gonEx4 [Pmyo-3::sfp-1]* | 19.13±0.60 | 19 | 15 | 75/86 | | 0.219 | 4A |  |
| *gonEx5 [Pregf-1::sfp-1]* | 18.17±0.55 | 18 | 15 | 78/88 | | 0.978 | 4A |  |
| N2 | 18.23±0.70 | 19 | 15 | 74/89 | | - | exp. 2 | 11-29-2021 |
| *gonEx17 [Pges-1::sfp-1]* | 13.69±0.39 | 13 | 11 | 88/97 | | <0.0001 |  |  |
| *gonEx18 [Psmu-1::sfp-1]* | 17.44±0.61 | 18 | 15 | 43/80 | | 0.241 |  |  |
| *gonEx4 [Pmyo-3::sfp-1]* | 18.91±0.60 | 19 | 15 | 75/86 | | 0.941 |  |  |
| *gonEx5 [Pregf-1::sfp-1]* | 18.22±0.46 | 18 | 14 | 80/98 | | 0.192 |  |  |
| N2 | 18.04±0.49 | 18 | 15 | 79/100 | | - | 4E | 06-05-2022 |
| *gonEx17 [Pges-1::sfp-1]* | 13.97±0.49 | 13 | 11 | 84/91 | | <0.0001 | 4E |  |
| *gonEx19 [Pges-1::△NTF2L]* | 17.30±0.55 | 17 | 13 | 70/90 | | 0.390 | 4E |  |
| *gonEx20 [Pges-1::△98-132]* | 17.48±0.84 | 16 | 14 | 46/67 | | 0.857 | 4E |  |
| *gonEx21 [Pges-1::△signal peptide]* | 15.20±0.47 | 14 | 12 | 82/100 | | <0.0001 | 4E |  |
| N2 | 17.87±0.57 | 18 | 15 | 82/102 | | - | exp. 2 | 07-19-2021 |
| *gonEx17 [Pges-1::sfp-1]* | 14.19±0.46 | 13 | 11 | 48/56 | | <0.0001 |  |  |
| *gonEx19 [Pges-1::△NTF2L]* | 17.34±0.54 | 17 | 14 | 70/90 | | 0.256 |  |  |
| *gonEx20 [Pges-1::△98-132]* | 15.71±0.78 | 16 | 12 | 75/86 | | 0.045 |  |  |
| *gonEx21 [Pges-1::△signal peptide]* | 16.16±0.46 | 16 | 14 | 73/99 | | 0.004 |  |  |
| N2 | 17.40±0.79 | 18 | 14 | 48/53 | | - | S4E | 03-09-2024 |
| *gonEx6 [Pges-1::K12H6.5]* | 19.61±0.46 | 20 | 16 | 77/86 | | 0.058 |  |  |
| *gonEx7 [Pges-1::F40G9.15]* | 16.74±0.53 | 17 | 14 | 57/70 | | 0.194 |  |  |
| N2 x N2♂ | 9.33±0.49 | 8 | 7 | 25/27 | | - | 5A | 01-01-2024 |
| N2 x *sfp-1*♂ | 12.60±0.72 | 12 | 10 | 21/28 | | <0.0001 | 5A |  |
| *daf-16（mgDf47）*x N2♂ | 8.55±0.36 | 9 | 7 | 12/22 | | 0.364 | 5A |  |
| *daf-16（mgDf47）*x *sfp-1*♂ | 11.08±0.55 | 11 | 9 | 21/31 | | 0.017 | 5A |  |
| N2 x N2♂ | 9.33±0.49 | 8 | 7 | 25/27 | | - | exp. 2 | 01-06-2024 |
| N2 x *sfp-1*♂ | 12.84±0.73 | 13 | 10 | 22/29 | | <0.0001 |  |  |
| *daf-16（mgDf47）*x N2♂ | 9.68±0.44 | 9 | 8 | 20/23 | | 0.574 |  |  |
| *daf-16（mgDf47）*x *sfp-1*♂ | 11.11±0.64 | 11 | 8 | 27/30 | | 0.024 |  |  |
| N2 x N2♂ | 10.39±0.48 | 10 | 9 | 24/26 | | - | 5B | 03-17-2024 |
| N2 x *sfp-1*♂ | 12.16±0.69 | 12 | 10 | 17/23 | | 0.031 | 5B |  |
| *skn-1（zu135）*x N2♂ | 11.00±0.45 | 11 | 9 | 26/30 | | 0.426 | 5B |  |
| *skn-1（zu135）*x *sfp-1*♂ | 10.74±0.51 | 11 | 9 | 19/24 | | 0.739 | 5B |  |
| N2 x N2♂ | 10.66±0.50 | 11 | 10 | 21/27 | | - | exp. 2 | 03-06-2024 |
| N2 x *sfp-1*♂ | 14.63±0.71 | 15 | 13 | 21/24 | | <0.0001 |  |  |
| *skn-1（zu135）*x N2♂ | 10.48±0.58 | 11 | 9 | 24/26 | | 0.899 |  |  |
| *skn-1（zu135）*x *sfp-1*♂ | 10.79±0.78 | 11 | 8 | 21/30 | | 0.394 |  |  |
| N2 x N2♂ | 10.00±0.80 | 10 | 7 | 22/28 | | - | exp. 3 | 03-16-2024 |
| N2 x *sfp-1*♂ | 14.31±0.64 | 14 | 12 | 16/18 | | 0.001 |  |  |
| *skn-1（zu135）*x N2♂ | 10.11±0.65 | 10 | 8 | 18/25 | | 0.936 |  |  |
| *skn-1（zu135）*x *sfp-1*♂ | 11.31±0.41 | 11 | 9 | 15/22 | | 0.628 |  |  |
| N2 | 17.01±0.45 | 17 | 13 | 119/136 | | - | 5E | 02-06-2024 |
| *gonEx17 [Pges-1::sfp-1]* | 14.33±0.50 | 14 | 12 | 48/52 | | <0.0001 | 5E |  |
| *skn-1（zu135）* | 12.40±0.50 | 12 | 10 | 57/63 | | <0.0001 | 5E |  |
| *gonEx17 [Pges-1::sfp-1];* *skn-1（zu135）* | 16.08±0.50 | 16 | 13 | 51/54 | | 0.026 | 5E |  |
| N2 | 16.79±0.58 | 17 | 13 | 70/80 | | - | exp. 2 | 02-26-2024 |
| *gonEx17 [Pges-1::sfp-1]* | 14.21±0.51 | 14 | 12 | 46/50 | | <0.0001 |  |  |
| *skn-1（zu135）* | 10.45±0.67 | 10 | 7 | 29/42 | | <0.0001 |  |  |
| *gonEx17 [Pges-1::sfp-1];* *skn-1（zu135）* | 13.19±0.55 | 12 | 10 | 62/73 | | <0.0001 |  |  |
| N2 | 16.75±0.76 | 15 | 13 | 53/58 | | - | exp. 3 | 03-13-2024 |
| *gonEx17 [Pges-1::sfp-1]* | 12.96±0.40 | 12 | 10 | 81/87 | | <0.0001 |  |  |
| *skn-1（zu135）* | 12.41±0.45 | 12 | 10 | 46/48 | | <0.0001 |  |  |
| *gonEx17 [Pges-1::sfp-1];* *skn-1（zu135）* | 12.62±0.46 | 12 | 10 | 55/63 | | <0.0001 |  |  |
| *gonEx17*-vector RNAi | 16.76±0.51 | 16 | 13 | 70/74 | | - | S5A | 02-18-2023 |
| *gonEx17-pqm-1* RNAi | 17.77±0.80 | 17 | 14 | 52/62 | | 0.238 | S5A |  |
| *gonEx17*-*ceh-60* RNAi | 14.58±0.52 | 15 | 12 | 55/60 | | 0.002 | S5A |  |
| *gonEx17-pha-4* RNAi | 15.33±0.62 | 13 | 12 | 57/60 | | 0.316 | S5A |  |
| *gonEx17*-*hsf-1* RNAi | 12.43±0.40 | 13 | 10 | 51/58 | | <0.0001 | S5A |  |
| *gonEx17-daf-12* RNAi | 15.26±0.52 | 14 | 12 | 74/77 | | 0.069 | S5A |  |
| *gonEx17*-vector RNAi | 15.81±0.58 | 16 | 12 | 58/64 | | - | exp. 2 | 12-17-2022 |
| *gonEx17-pqm-1* RNAi | 17.02±0.61 | 17 | 13 | 45/52 | | 0.227 |  |  |
| *gonEx17*-ceh-60 RNAi | 15.00±0.71 | 14 | 12 | 31/40 | 0.771 | |  |  |
| N2-vector RNAi | 20.71±0.55 | 21 | 16 | 92/100 | - | | S5B | 02-18-2023 |
| N2*-pqm-1* RNAi | 15.16±0.56 | 14 | 11 | 88/94 | <0.0001 | | S5B |  |
| N2-*ceh-60* RNAi  N2-*pha-4* RNAi | 21.19±0.79  20.36±0.53 | 22  20 | 16  17 | 48/49  80/82 | 0.601  0.258 | | S5B  S5B |  |
| N2-*hsf-1* RNAi | 11.66±0.31 | 12 | 10 | 56/64 | <0.0001 | | S5B |  |
| N2*-daf-12* RNAi | 19.44±0.49 | 19 | 16 | 80/92 | 0.007 | | S5B |  |
| N2-vector RNAi | 21.60±0.70 | 21 | 16 | 56/62 | - | | exp. 2 | 12-17-2022 |
| N2*-pqm-1* RNAi | 16.22±0.56 | 15 | 12 | 56/60 | <0.0001 | |  |  |
| N2-*ceh-60* RNAi | 21.97±0.94 | 22 | 16 | 38/40 | 0.449 | |  |  |
| N2-vector RNAi | 22.00±0.43 | 22 | 19 | 116/135 | - | | S5E | 11-13-2023 |
| N2*-sek-1* RNAi | 19.81±0.64 | 20 | 15 | 71/81 | 0.021 | | S5E |  |
| *gonEx17*-vector RNAi | 13.43±0.71 | 14 | 11 | 37/38 | <0.0001 | | S5E |  |
| *gonEx17-sek-1* RNAi | 15.71±0.48 | 16 | 13 | 66/72 | <0.0001 | | S5E |  |
| N2-vector RNAi | 22.00±0.43 | 22 | 19 | 116/135 | - | | S5F | 10-20-2023 |
| N2*-pmk-1* RNAi | 19.79±0.58 | 20 | 16 | 42/47 | 0.001 | | S5F |  |
| *gonEx17*-vector RNAi | 13.43±0.71 | 14 | 11 | 37/38 | <0.0001 | | S5F |  |
| *gonEx17-pmk-1* RNAi | 16.18±0.52 | 16 | 14 | 57/58 | <0.0001 | | S5F |  |
| N2-vector RNAi | 21.83±0.55 | 22 | 19 | 84/95 | - | | exp. 2 | 10-30-2023 |
| N2*-pmk-1* RNAi | 17.77±0.43 | 18 | 15 | 42/47 | <0.0001 | |  |  |
| *gonEx17*-vector RNAi | 14.73±0.43 | 14 | 11 | 64/70 | <0.0001 | |  |  |
| *gonEx17-pmk-1* RNAi | 16.20±0.45 | 16 | 13 | 43/44 | <0.0001 | |  |  |
| *N2-vector RNAi* | 22.39±0.55 | 23 | 19 | 68/85 | - | | exp. 3 | 11-30-2023 |
| *N2-pmk-1 RNAi* | 20.10±0.55 | 21 | 17 | 67/71 | <0.0001 | |  |  |
| *gonEx17-vector RNAi* | 14.98±0.46 | 15 | 11 | 71/80 | <0.0001 | |  |  |
| *gonEx17-pmk-1 RNA* | 15.66±0.54 | 14 | 12 | 75/82 | <0.0001 | |  |  |
| N2-vector RNAi | 22.79±0.61 | 24 | 20 | 49/50 | - | | S6F | 11-06-2023 |
| N2*-vit-1* RNAi | 17.81±0.71 | 18 | 11 | 63/66 | <0.0001 | | S6F |  |
| *gonEx17*-vector RNAi | 15.33±0.42 | 15 | 12 | 67/86 | <0.0001 | | S6F |  |
| *gonEx17-vit-1* RNAi | 13.84±0.42 | 13 | 11 | 78/82 | <0.0001 | | S6F |  |
| N2-vector RNAi | 22.79±0.61 | 24 | 20 | 49/50 | - | | S6G | 11-08-2023 |
| N2*-vit-2* RNAi | 18.15±0.73 | 18 | 13 | 48/59 | <0.0001 | | S6G |  |
| *gonEx17*-vector RNAi | 15.33±0.42 | 15 | 12 | 67/86 | <0.0001 | | S6G |  |
| *gonEx17-vit-2* RNAi | 14.72±0.62 | 14 | 10 | 59/61 | <0.0001 | | S6G |  |
| N2-vector RNAi | 22.79±0.61 | 24 | 20 | 49/50 | - | | S6H | 11-10-2023 |
| N2*-vit-3* RNAi | 18.06±0.53 | 19 | 13 | 88/93 | <0.0001 | | S6H |  |
| *gonEx17*-vector RNAi | 15.33±0.42 | 15 | 12 | 67/86 | <0.0001 | | S6H |  |
| *gonEx17-vit-3* RNAi | 13.89±0.38 | 13 | 11 | 91/100 | <0.0001 | | S6H |  |
| N2-vector RNAi | 22.79±0.61 | 24 | 20 | 49/50 | - | | S6I | 11-15-2023 |
| N2*-vit-4* RNAi | 16.71±0.66 | 17 | 13 | 40/42 | <0.0001 | | S6I |  |
| *gonEx17*-vector RNAi | 15.33±0.42 | 15 | 12 | 67/86 | <0.0001 | | S6I |  |
| *gonEx17-vit-4* RNAi | 13.40±0.45 | 13 | 10 | 62/74 | <0.0001 | | S6I |  |
| N2-vector RNAi | 22.79±0.61 | 24 | 20 | 49/50 | - | | S6J | 11-15-2023 |
| N2*-vit-5* RNAi | 17.70±0.51 | 17 | 13 | 105/119 | <0.0001 | | S6J |  |
| *gonEx17*-vector RNAi | 15.33±0.42 | 15 | 12 | 67/86 | <0.0001 | | S6J |  |
| *gonEx17-vit-5* RNAi | 12.73±0.35 | 13 | 10 | 84/102 | <0.0001 | | S6J |  |
| N2-vector RNAi | 22.79±0.61 | 24 | 20 | 49/50 | - | | S6K | 11-15-2023 |
| N2*-vit-6* RNAi | 17.50±0.59 | 17 | 12 | 78/90 | <0.0001 | | S6K |  |
| *gonEx17*-vector RNAi | 15.33±0.42 | 15 | 12 | 67/86 | <0.0001 | | S6K |  |
| *gonEx17-vit-6* RNAi | 14.31±0.59 | 13 | 11 | 54/70 | <0.0001 | | S6K |  |
| N2-vector RNAi | 22.41±0.75 | 22 | 18 | 64/95 | - | | 7E | 09-30-2022 |
| N2*-fat-1* RNAi | 21.90±0.55 | 23 | 17 | 85/99 | 0.202 | | 7E |  |
| *gonEx17*-vector RNAi | 18.39±0.49 | 18 | 15 | 77/96 | <0.0001 | | 7E |  |
| *gonEx17-fat-1* RNAi | 24.00±0.54 | 25 | 21 | 93/102 | 0.479 | | 7E |  |
| N2-vector RNAi | 20.81±1.01 | 20 | 16 | 36/43 | - | | exp. 2 | 11-20-2022 |
| N2*-fat-1* RNAi | 22.11±0.81 | 23 | 16 | 53/81 | 0.585 | |  |  |
| *gonEx17*-vector RNAi | 17.29±0.41 | 17 | 13 | 113/121 | <0.0001 | |  |  |
| *gonEx17-fat-1* RNAi | 19.79±1.00 | 19 | 14 | 42/50 | 0.475 | |  |  |
| N2-vector RNAi | 20.81±1.01 | 20 | 16 | 36/43 | - | | 7F | 11-20-2022 |
| N2*-fat-2* RNAi | 21.56±0.76 | 23 | 17 | 59/68 | 0.905 | | 7F |  |
| *gonEx17*-vector RNAi | 17.29±0.41 | 17 | 13 | 113/121 | <0.0001 | | 7F |  |
| *gonEx17-fat-2* RNAi | 22.03±0.76 | 24 | 18 | 60/67 | 0.661 | | 7F |  |
| N2-vector RNAi | 22.41±0.75 | 22 | 18 | 64/95 | - | | exp. 2 | 09-30-2022 |
| N2*-fat-2* RNAi | 17.31±0.94 | 15 | 14 | 45/73 | <0.0001 | |  |  |
| *gonEx17*-vector RNAi | 18.39±0.49 | 18 | 15 | 77/96 | <0.0001 | |  |  |
| *gonEx17-fat-2* RNAi | 21.23±0.89 | 20 | 15 | 70/105 | 0.792 | |  |  |
| N2-vector RNAi | 22.41±0.75 | 22 | 18 | 64/95 | - | | S7A | 09-30-2022 |
| N2*-fat-3* RNAi | 23.96±0.52 | 26 | 20 | 138/142 | 0.241 | | S7A |  |
| *gonEx17*-vector RNAi | 18.39±0.49 | 18 | 15 | 77/96 | <0.0001 | | S7A |  |
| *gonEx17-fat-3* RNAi | 23.32±0.62 | 24 | 18 | 98/109 | 0.422 | | S7A |  |
| N2-vector RNAi | 22.41±0.75 | 22 | 18 | 64/95 | - | | S7B | 09-30-2022 |
| N2*-fat-4* RNAi | 19.91±0.60 | 21 | 17 | 74/90 | 0.002 | | S7B |  |
| *gonEx17*-vector RNAi | 18.39±0.49 | 18 | 15 | 77/96 | <0.0001 | | S7B |  |
| *gonEx17-fat-4* RNAi | 22.15±0.75 | 23 | 17 | 93/94 | 0.554 | | S7B |  |
| N2-vector RNAi | 20.81±1.01 | 20 | 16 | 36/43 | - | | S7C | 11-20-2022 |
| N2*-fat-5* RNAi | 21.31±0.58 | 22 | 18 | 84/98 | 0.838 | | S7C |  |
| *gonEx17*-vector RNAi | 17.29±0.41 | 17 | 13 | 113/121 | <0.0001 | | S7C |  |
| *gonEx17-fat-5* RNAi | 21.06±0.94 | 23 | 16 | 48/68 | 0.915 | | S7C |  |
| N2-vector RNAi | 22.41±0.75 | 22 | 18 | 64/95 | - | | exp. 2 | 09-30-2022 |
| N2-*fat-5* RNAi | 22.35±0.66 | 22 | 18 | 84/94 | 0.064 | |  |  |
| gonEx17-vector RNAi | 18.39±0.49 | 18 | 15 | 77/96 | <0.0001 | |  |  |
| gonEx17-*fat-5* RNAi | 22.99±0.82 | 23 | 17 | 58/70 | 0.821 | |  |  |
| *N2-vector RNAi* | 20.81±1.01 | 20 | 16 | 36/43 | - | | S7D | 11-20-2022 |
| *N2-fat-6 RNAi* | 13.71±0.43 | 13 | 12 | 61/61 | <0.0001 | | S7D |  |
| *gonEx17-vector RNAi* | 17.29±0.41 | 17 | 13 | 113/121 | <0.0001 | | S7D |  |
| *gonEx17-fat-6 RNAi* | 14.89±0.42 | 15 | 12 | 84/97 | <0.0001 | | S7D |  |
| *N2-vector RNAi* | 22.41±0.75 | 22 | 18 | 64/95 | - | | exp. 2 | 09-30-2022 |
| *N2-fat-6 RNAi* | 14.64±0.43 | 14 | 13 | 64/99 | <0.0001 | |  |  |
| *gonEx17-vector RNAi* | 18.39±0.49 | 18 | 15 | 77/96 | <0.0001 | |  |  |
| *gonEx17-fat-6 RNAi* | 15.32±0.49 | 15 | 13 | 55/92 | <0.0001 | |  |  |
| *N2-vector RNAi* | 22.41±0.75 | 22 | 18 | 64/95 | - | | S7E | 09-30-2022 |
| *N2-fat-7 RNAi* | 14.64±0.43 | 14 | 13 | 64/99 | <0.0001 | | S7E |  |
| *gonEx17-vector RNAi* | 18.39±0.49 | 18 | 15 | 77/96 | <0.0001 | | S7E |  |
| *gonEx17-fat-7 RNAi* | 15.32±0.49 | 15 | 13 | 55/92 | <0.0001 | | S7E |  |
| *N2-vector RNAi* | 20.81±1.01 | 20 | 16 | 36/43 | - | | exp. 2 | 11-20-2022 |
| *N2-fat-7 RNAi* | 16.90±0.46 | 17 | 14 | 63/78 | <0.0001 | |  |  |
| *gonEx17-vector RNAi* | 17.29±0.41 | 17 | 13 | 113/121 | <0.0001 | |  |  |
| *gonEx17-fat-7 RNAi* | 17.45±0.52 | 17 | 14 | 44/49 | <0.0001 | |  |  |
